# DNA-PKcs governs LAT-dependent signaling in CD4^+^ and CD8^+^ T cells

**DOI:** 10.1101/2025.03.06.641745

**Authors:** Randall R. Rainwater, Ana C. Azevedo-Pouly, Zachary J. Waldrip, Belle H. Hicks, Nicholas A. Callais, Brian Koss, Aaron J. Storey, Lyle Burdine, Marie Schluterman Burdine

## Abstract

Formation of the immune synapse (IS) following T cell antigen recognition includes recruitment of the Linker for Activation of T cells (LAT). Once at the IS, LAT tyrosines are phosphorylated allowing it to serve as a scaffold for formation of the “signalosome”, a multiprotein complex that drives TCR signaling. Here, we show that upon T cell activation, DNA dependent protein kinase catalytic subunit (DNA-PKcs) interacts with LAT and localizes to the IS. Inhibition of DNA-PKcs diminishes LAT localization at the IS. We identified two LAT serines phosphorylated by DNA-PKcs, S224 and S241, that impact LAT tyrosine phosphorylation, protein binding, and cytokine production. Using our mouse model designed to delete DNA-PKcs expression within mature CD4^+^ or CD8^+^ T cells, we show loss of DNA-PKcs results in T cells unable to control tumor growth or induce allogeneic graft rejection. These data bring to the forefront DNA-PKcs as a pivotal protein in T cell function.

## Introduction

Antigen ligation of the T cell receptor (TCR) triggers the formation of a supramolecular structure known as the immune synapse (IS) at the interface between the TCR and the antigen-presenting cell (APC).^1^ This involves polarization of the T cell toward the APC and reorganization of cytoskeletal elements and organelles, including the Golgi complex, beneath the IS. This facilitates delivery of critical signaling molecules to the IS to initiate molecular pathways impacting T cell response, including transcriptional pathways for differentiation into antigen-specific effector cells. Pharmaceutical targeting of IS proteins led to the development of effective therapies for cancer and infectious diseases as well as promising approaches for preventing transplant rejection.^2^ It is evident that continuing to explore the IS landscape to identify proteins involved in regulating T cell signaling can yield significant therapeutic benefits.

A critical step in IS signal transduction is recruitment of the transmembrane adaptor molecule Linker for Activated T cells (LAT) to the synaptic membrane. While many questions remain regarding the trafficking of LAT to the IS, it is well documented that upon TCR/antigen binding, LAT translocates to the IS where it is phosphorylated by the tyrosine kinase ZAP70.^3–5^ The phosphorylated tyrosine’s serve as binding sites for signaling molecules forming the multimolecular complex known as the “LAT signalosome”.^6^ This complex brings effector molecules into contact to drive proximal TCR signaling and the subsequent T cell response. Loss of LAT expression or mutation of key tyrosine phosphorylation sites results in the misregulation of T cells in animal models.^7–10^

In its canonical role, the phosphoinositide 3-kinase (PI3K)-related kinase DNA-dependent protein kinase catalytic subunit (DNA-PKcs), along with family members ATM and ATR, functions in the nucleus as a double-strand DNA damage repair (DDR) kinase to promote nonhomologous end joining (NHEJ).^11,12^ It was first linked to the developing immune system when mice with a severe combined immunodeficiency (SCID) phenotype, characterized by deficiencies in mature T and B cells, were discovered to harbor DNA-PKcs loss-of-function mutations.^13–16^ This defect is attributed to DNA-PKcs’s role in V(D)J recombination, which promotes B and T cell receptor diversity.

Though less well characterized, DNA-PKcs also functions in mature immune cells independent of its role in DNA damage repair. For instance, DNA-PKcs is required for Toll-like receptor 9 (TLR9) signaling and interferon (IFN)-α/β production in dendritic cells, functioning as a viral DNA sensor and supporting efficient antigen presentation to drive a Th2-mediated response.^17,18^ DNA-PKcs also regulates differentiation of CD4⁺ T cells into Th1 and Th2 subtypes by altering the expression of Gata3 and T-bet.^19^ Additionally, DNA-PKcs, in conjunction with the KU complex expressed in the cytoplasm of aging CD4⁺ T cells, acts as a sensor for accumulated cytoplasmic DNA, resulting in heightened T cell activation.^20^ Here, we provide further evidence that DNA-PKcs is a critical component of the adaptive immune response and suggest a new mechanism by which it impacts CD4⁺ and CD8⁺ T cell function. For the first time, we demonstrate that upon T cell activation, DNA-PKcs physically interacts with LAT and is required for LAT localization to the IS. We identified two serines within LAT, S224 and S241, that are phosphorylated by DNA-PKcs and are critical for LAT tyrosine phosphorylation and IL2 and IFN*γ* production. Using unique transgenic mouse models designed to delete DNA-PKcs expression specifically within mature CD4⁺ or CD8⁺ T cells, we show that loss of DNA-PKcs results in T cells unable to control tumor growth or induce allogeneic transplant graft rejection. These data highlight a new mechanism influencing T cell responses with potential therapeutic application for immune-related diseases.

## Results

### T cell activation uniquely activates DNA-PKcs in the cytoplasm

The function of DNA-PKcs within the cytoplasm of T cells remains poorly understood. Using cell fractionation and laser confocal microscopy (LSCM), we demonstrate that upon activation of human Jurkat T cells with αCD3/CD28, active DNA-PKcs indicated by the presence of phosphorylated serine 2056 (pDNA-PKcs) is specifically elevated within the cytoplasm (**Figures 1A, B, C**).^21,22^ Western blot analysis of cytosolic fractions revealed that when T cells were stimulated with αCD3/CD28 for five minutes and then removed from the stimulus, pDNA-PKcs levels peaked at 15 minutes and returned to baseline by 45 minutes (**Figures S1A, B**).

**Figure 1.**
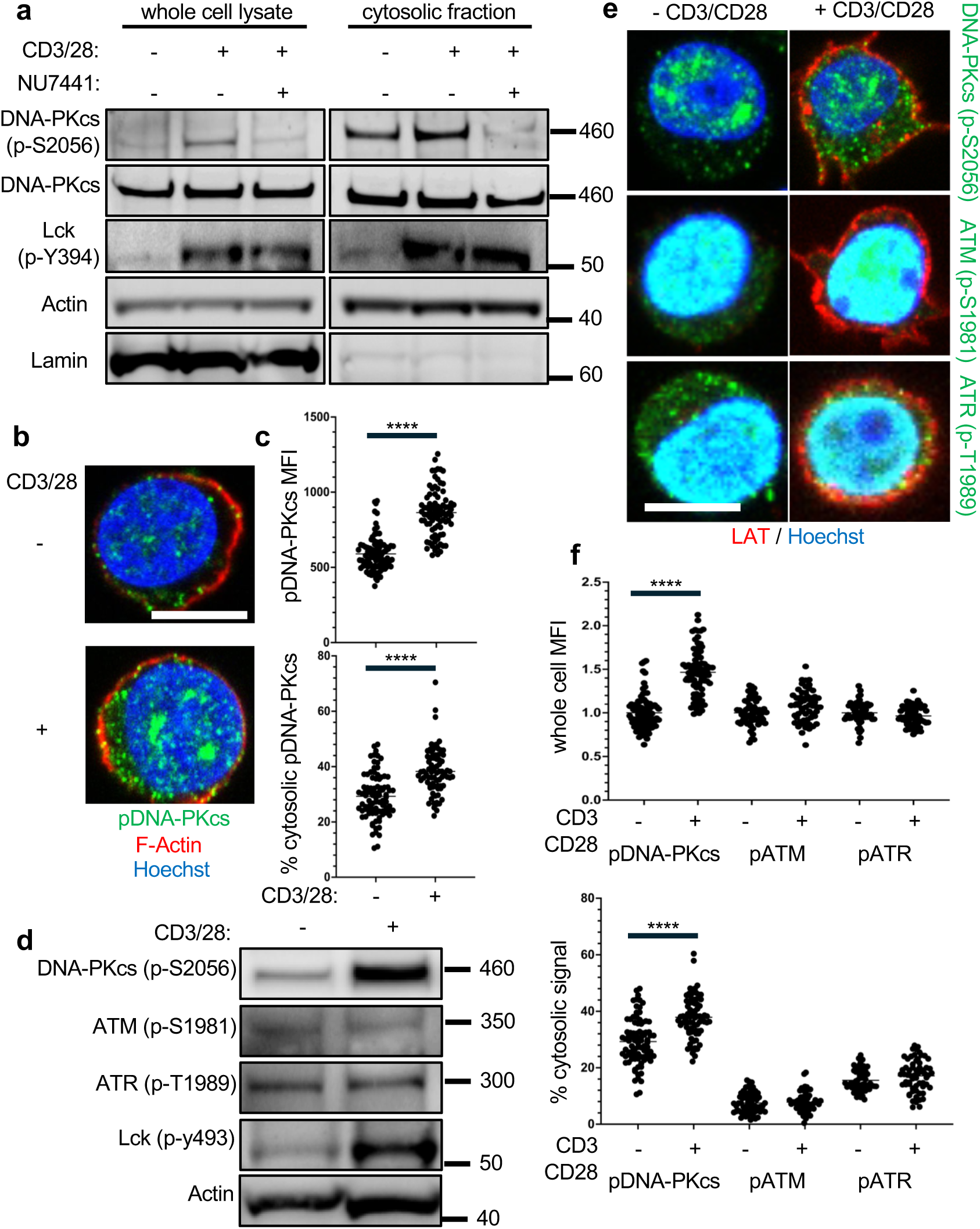
DNA-PKcs expression increases in the cytoplasm and at the plasma membrane following T-cell activation. **A)** Subcellular fractionation of E6.1 Jurkat T cells shows increased DNA-PKcs phosphorylation at S2056 after 2 minutes of αCD3/CD28 TCR stimulation, which is reduced by the DNA-PKcs inhibitor NU7441 (5 µM). **B)** LSCM imaging of Jurkat T cells revealed that two minutes of αCD3/CD28 TCR stimulation led to increased protein expression (green) in the cytosol and at the plasma membrane alongside F-Actin (red). Quantification of pDNA-PKcs by mean fluorescent intensity (MFI) is shown for **C)** whole-cell pDNA-PKcs and the cytosolic ratio of pDNA-PKcs. **D, E)** Among PIKK family members, only DNA-PKcs, but not ATM or ATR, is activated in the cytosol following TCR stimulation. **F)** MFI values and the cytosolic-to-whole-cell ratio of phosphorylated PIKKs. Scale bar = 5 µm. Representative western blots and images from n=3 independent experiments. Statistical significance was determined using unpaired T-test (α=0.05, ****p<0.0001).

Interestingly, a portion of cytoplasmic pDNA-PKcs localized to the plasma membrane upon αCD3/CD28 activation (**Figures 1B, C**). We investigated whether this phenomenon was unique to DNA-PKcs or if related DDR kinases ATM and ATR exhibited similar expression patterns following T cell activation.^23^ After stimulation of Jurkat T cells, we performed western blot analysis on cytosolic fractions to assess the levels of activated ATM (pS1981) and ATR (pS1989). Unlike DNA-PKcs, TCR stimulation did not induce an increase in pATM or pATR in the cytoplasm. This finding was corroborated by LSCM results, which showed that while pATM and pATR levels increased overall, there was not an increase in the cytosol as seen with pDNA-PKcs (**Figures 1D, E, F**). These results suggest DNA-PKcs plays a unique role in T cells, distinct from its well-known function in nuclear DNA damage response (DDR) and point to an unexplored step in TCR signaling that involves DNA-PKcs’s localization at the plasma membrane.

### LAT is a phosphorylation target of DNA-PKcs

To define the function of DNA-PKcs at the plasma membrane of activated T cells, we evaluated our recently published phospho-mass spectrometry study, which analyzed differentially phosphorylated proteins in activated Jurkat T cells treated with or without the DNA-PKcs inhibitor NU7441.^24,25^ We specifically focused on proteins known to localize to the plasma membrane of activated T cells. Notably, we identified the IS-associated protein LAT. Phosphorylation of LAT at serine 241 (S241) was significantly reduced (fold change >10) upon NU7441 treatment, indicating that DNA-PKcs phosphorylates LAT at this residue. We confirmed that DNA-PKcs phosphorylates LAT using an *in vitro* DNA-PKcs luminescent kinase assay and a LAT fragment containing the S241 amino acid (**Figures S2A, B**). Analysis of the phosphorylated LAT peptide by mass spectrometry confirmed that DNA-PKcs not only phosphorylates S241 but also serine 224 (S224). **Figure S2C** illustrates the ratios of phosphorylated S224 and S241 to unphosphorylated LAT in the presence and absence of DNA-PKcs. Sequence alignment of these phosphorylation sites reveals a high degree of conservation from mice to humans. Both serines are located within conserved S/Q motifs, known DNA-PKcs consensus sites (**Figure S2D**).

### DNA-PKcs inhibition diminishes LAT localization to the immune synapse

To further explore a functional relationship between DNA-PKcs and LAT, we analyzed expression patterns of LAT and DNA-PKcs by LSCM in Jurkat T cells following activation. As expected, activation resulted in clear localization of LAT to the plasma membrane. Interestingly, we observed co-localization between DNA-PKcs and LAT on the plasma membrane of activated T cells (**Figure 2A**). LAT is found specifically localized to the IS following antigen recognition. To form a true, “visible” IS, we cocultured Jurkat T cells with Raji B cells preloaded with the Staphylococcal enterotoxin type E (SEE) antigen.^26^ LSCM images in **Figure 2B** show LAT and pDNA-PKcs co-localized to the IS at the T-B cell interface. Moreover, we show that DNA-PKcs physically interacts with LAT as DNA-PKcs co-immunoprecipitated with the pull-down of LAT (**Figure 2C**).

**Figure 2.**
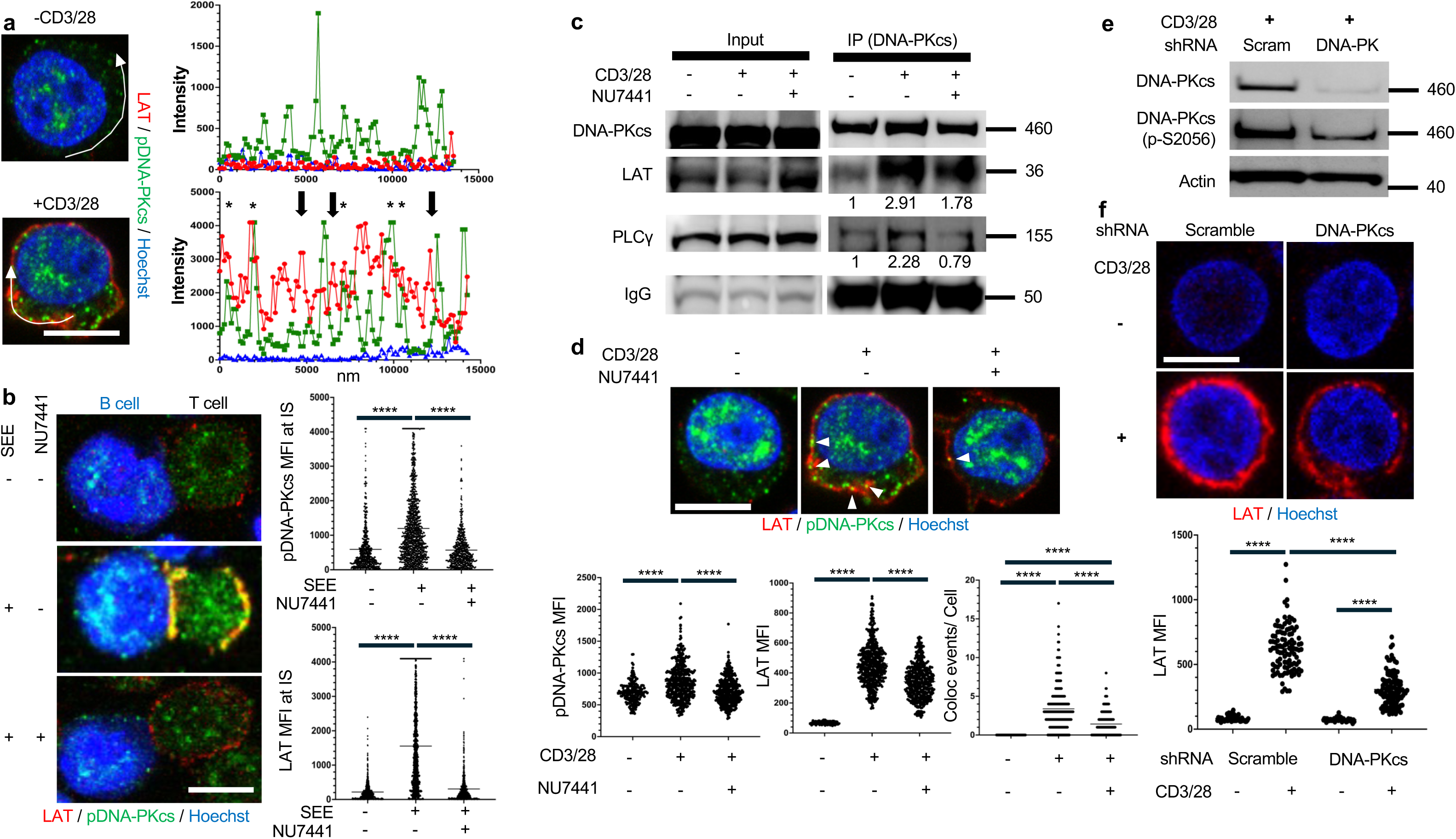
DNA-PKcs inhibition reduces LAT localization at the immune synapse. **A)** Histogram intensity analysis of 2-minute αCD3/CD28 TCR stimulated Jurkat T cells highlights areas of colocalization where pDNA-PKcs and LAT peaks overlap (*) and areas of single molecule expression where peaks do not overlap (arrow). **B)** In SEE-pulsed Raji B cells cultured with Jurkat T cells, pDNA-PKcs (green) colocalizes with LAT (red) at the immune synapse (IS), quantified by mean fluorescent intensity (MFI). **C)** Co-immunoprecipitation analysis in Jurkat T cells shows DNA-PKcs interacts with LAT following TCR stimulation, which is reduced by the DNA-PKcs inhibitor NU7441. **D)** LSCM demonstrates LAT (red) localization at the plasma membrane with pDNA-PKcs (green) upon TCR stimulation, which decreases with NU7441 (5 µM). Quantification reveals significant increases in MFI for pDNA-PKcs and LAT after TCR stimulation, with decreases upon DNA-PKcs inhibition. **E)** shRNA-mediated knockdown of DNA-PKcs (>90% reduction) reduces total DNA-PKcs expression on western blot. **F)** shRNA-mediated knockdown of DNA-PKcs reduces LAT (red) localization at the plasma membrane after 2-minute αCD3/CD28 TCR stimulation. Scale bar = 5 µm. Representative images from n=3 independent experiments. Multiple comparisons ANOVA (α=0.05, ****p < 0.0001, ***p < 0.0005).

Surprisingly, inhibiting DNA-PKcs with NU7441, not only blocked DNA-PKcs localization but also abrogated expression of LAT at the IS and reduced the interaction between DNA-PKcs and LAT (**Figures 2B-D**). To further validate DNA-PKcs inhibition impacts LAT IS localization, we used shRNA methods to knockdown DNA-PKcs expression in Jurkat T cells and analyzed for LAT expression at the plasma membrane. Similar to our previous findings, loss of DNA-PKcs expression significantly reduced LAT expression at the plasma membrane (**Figure 2E**). This relationship between LAT and DNA-PKcs was further confirmed in primary human CD3^+^ T cells isolated from commercially-available human PBMCs. Activation with αCD3/CD28 stimulated the localization of LAT at the plasma membrane which was reduced with NU7441 treatment (**Figures S3A, B**). These novel findings indicate a mechanism by which DNA-PKcs is required for the translocation or retention of LAT at the IS.

### Phosphorylation at LAT S224 and S241 is not required for proliferation but is for IL2 production

We next investigated the relevance of the phosphorylation sites S224 and S241 to LAT activity and T cell function. To achieve this, we generated mCherry-tagged lentiviral expression vectors containing serine-to-alanine mutations at S224 and S241, designated as S224A and S241A, respectively, and a double mutant with both S224 and S241 mutated to alanines (S2A). These vectors, along with vectors expressing wild-type (WT) LAT, were transduced into the LAT-deficient Jurkat T cell line J.LAT, previously generated by the Arthur Weiss Laboratory.^27^ Following transduction, the expression of WT and mutant LAT was validated using flow cytometry for mCherry expression and western blot for LAT protein levels (**Figures S4A, B**).

First, we assessed whether loss of phosphorylation at these sites affects proliferation. J.LAT cells reconstituted with either WT LAT or the mutants (S224A, S241A, or S2A) expectedly proliferated at rates similar to the parental E6.1 Jurkat T cell line (**Figures S5A, B**) as loss of LAT in J.LAT cells was previously reported to not alter proliferation.^27^

Next, we evaluated the impact of the phosphorylation mutations on IL2 production. After transduction, T cells were activated with αCD3/CD28 with supernatant cytokine levels quantified using ELISA assays. While untransduced J.LAT cells were unable to produce measurable levels of the cytokine, J.LAT cells reconstituted with WT LAT produced significant amounts of IL2 following activation. In contrast, reconstitution with the S224A, S241A, or S2A mutants did not rescue IL2 production with levels comparable to untransduced J.LAT cells (**Figure 3A**). These data indicate that phosphorylation of serines S224 and S241 is crucial for regulating T cell signaling pathways.

**Figure 3.**
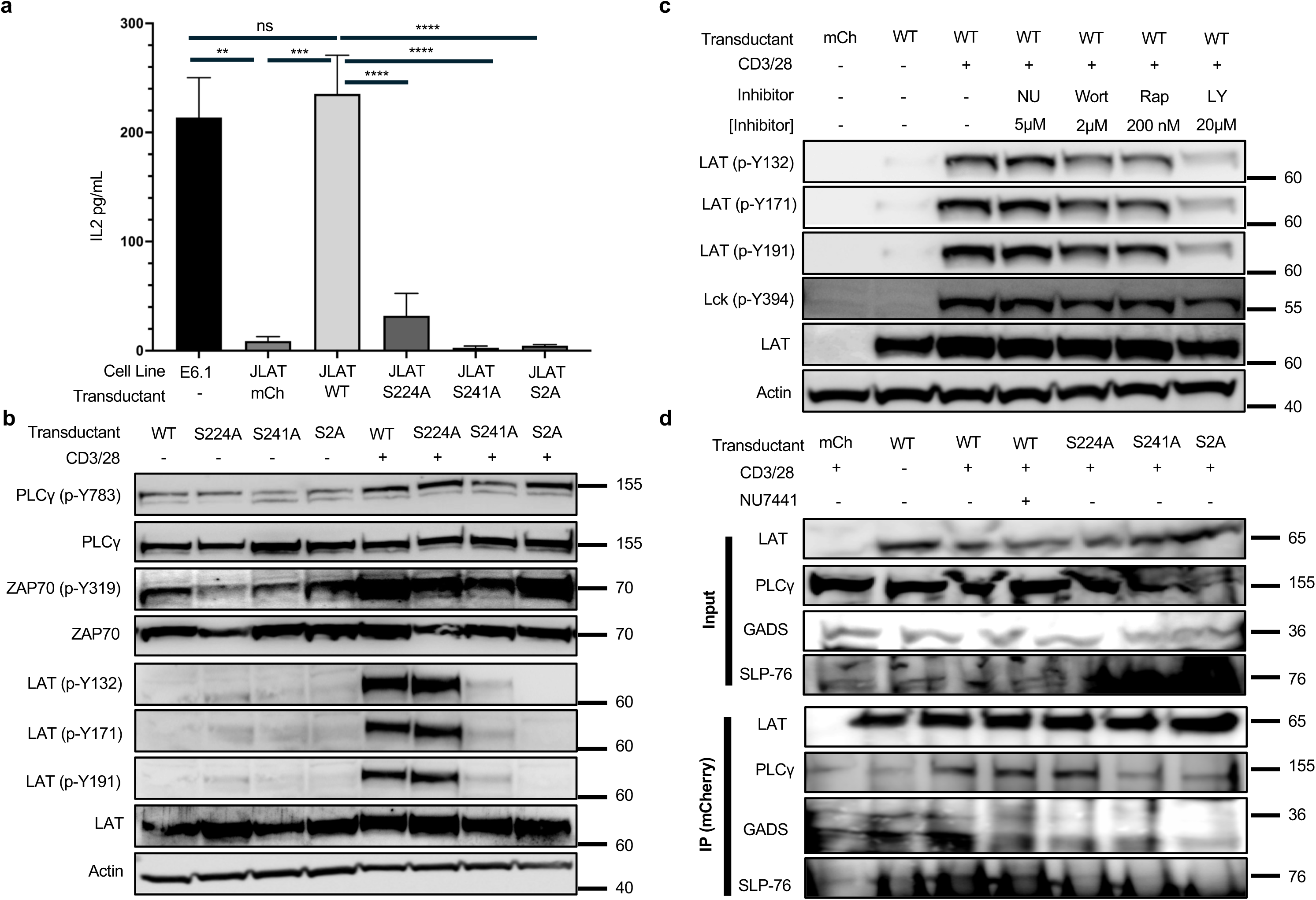
Loss of phosphorylation at S224 and S241 prevents LAT tyrosine phosphorylation, LAT protein binding, and cytokine production. **A)** JLAT transductants with S224 or S241 mutated are not able to produce IL2 with 24 hr αCD3/CD28 stimulation. **B)** Western blot analysis of tyrosine phosphorylation events on LAT (Y132, 171 and 191) in resting or 2-minute αCD3/CD28 TCR stimulated JLAT transductants. Following 2-minute αCD3/CD28 stimulation, Y132, Y171 and Y191 are phosphorylated in WT and S224A JLAT transductants but not in S241A and S2A JLATs. **C)** Phosphorylation of Y132, Y171 and Y191 of LAT are blocked with pan-PI3K inhibitor LY294002, but not selective PI3K inhibitors in WT LAT transduced JLAT T cells. **D)** Phosphorylation of LAT at S241 is required for signalosome formation given decreased pulldown of PLCγ, GADS, and SLP-76 in JLAT cells expressing S241A LAT and S2A LAT. n=3 independent experiment, multiple comparisons ANOVA (α=0.05, **p<0.01, ***p<0.0005, ****p< 0.0001).

### Phosphorylation at S224 and S241 impacts the LAT signalosome protein composition

Once LAT localizes to the IS, tyrosines in its cytoplasmic tail, specifically Y132, Y171, and Y191, undergo phosphorylation by the kinase ZAP70.^5^ These residues then serve as docking sites for signaling proteins that constitute the signalosome. The protein composition of the signalosome dictates downstream signaling pathways and ultimately determines how a T cell responds to an antigen.^6^ Analysis of these tyrosine phosphorylation sites following αCD3/CD28 stimulation revealed that phosphorylation at Y132, Y171, and Y191 was prevented in J.LAT cells reconstituted with either the S241A or S2A mutants, but not with S224A or WT LAT (**Figure 3B**). Interestingly, inhibition of DNA-PKcs with NU7441 did not block tyrosine phosphorylation, indicating other kinases are capable of phosphorylating these serines. This was supported by the observation that treatment of Jurkat T cells with the pan-PI3K inhibitor LY294002 blocked phosphorylation at Y132, Y171, and Y191 (**Figure 3C**).

Phosphorylation of these tyrosines is known to impact the binding of key LAT signalosome proteins, such as PLC*γ* and GADS (GRB2-related adaptor protein).^6,9^ Therefore, we investigated whether mutations at S224 or S241 hindered the binding of these proteins to LAT. Co-immunoprecipitation experiments performed in J.LAT cells reconstituted with WT LAT demonstrated binding between LAT and PLC*γ* and GADS following αCD3/CD28 activation, as expected. However, the S241A and S2A mutants were unable to interact with these proteins (**Figure 3D**). Since LAT indirectly binds to SLP-76 through its interaction with GADS, we also observed that the S241A and S2A mutants consequently prevented the interaction of SLP-76 with the LAT complex (**Figure 3D**).^28^ These findings indicate that phosphorylation of S241 is critical for the binding of proteins to LAT and regulates the composition of the signalosome.

To further elucidate how these phosphorylation sites affect the signalosome’s composition, we conducted a co-immunoprecipitation mass spectrometry study. This approach allowed us to comprehensively compare the binding patterns between WT LAT and the mutants. J.LAT cells reconstituted with either the mCherry vector alone, WT LAT, S224A LAT, or S241A LAT were activated with αCD3/CD28 for 15 minutes, followed by mCherry immunoprecipitation. Lysates were subjected to mass spectrometry. After filtering out non-specific binding proteins using the mCherry-only control samples and normalizing to LAT levels, we compared the binding patterns between activated WT and the S224A or S241A mutant cell lines. We identified significant changes in protein binding between WT and both mutants. Notably, the S241A mutant exhibited the most pronounced differences from WT (**Figure 4**). This observation is consistent with our findings that the S241A mutant showed defects in known tyrosine phosphorylation sites. However, further analysis of additional LAT tyrosine residues may reveal alterations in the S224A mutant that account for the observed changes in protein binding compared to WT.

**Figure 4.**
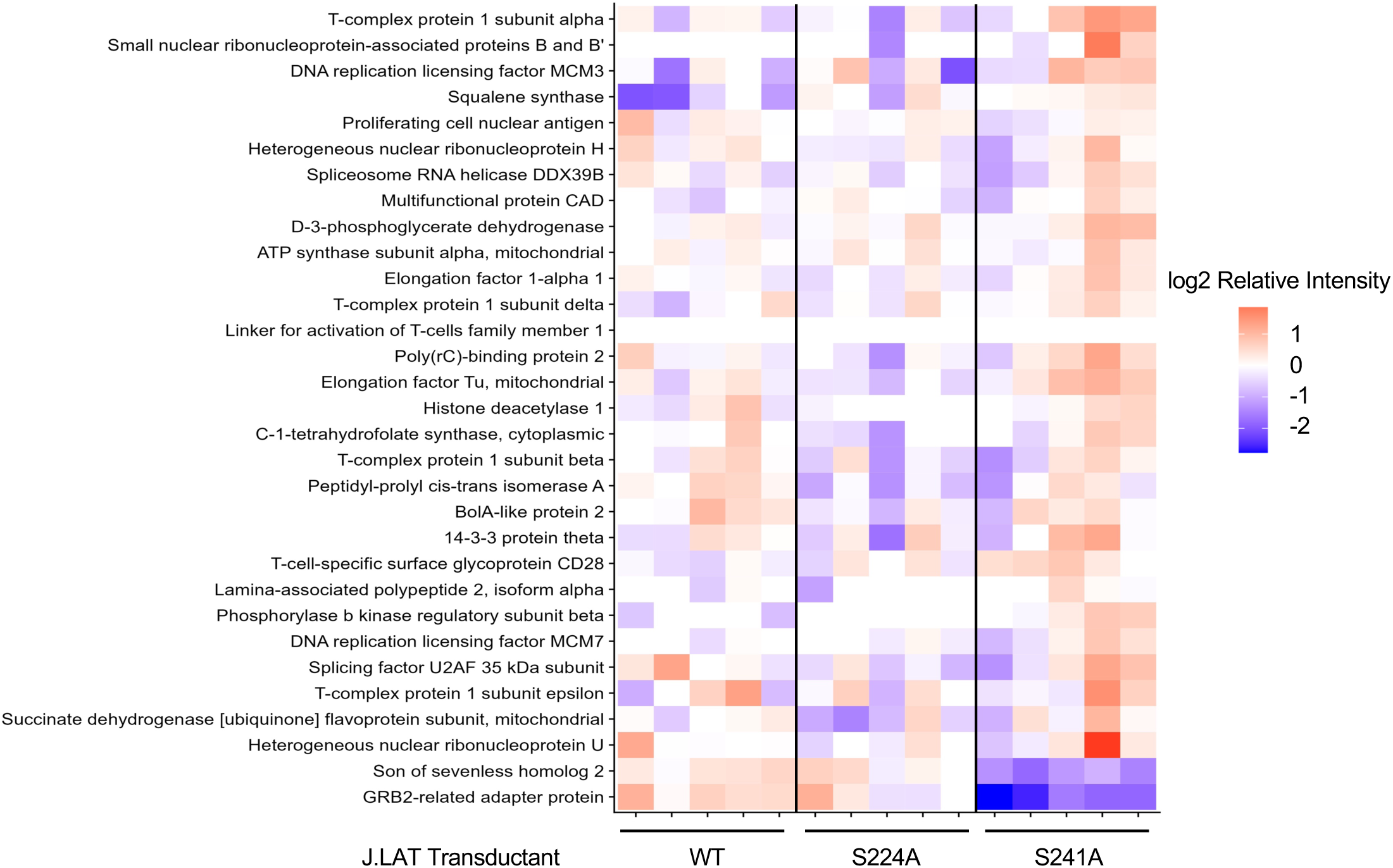
Loss of phosphorylation at S224 and S241 alters LAT signalosome formation. Co-IP coupled mass spec was completed in 15-minute αCD3/CD28 TCR stimulated JLAT transductants expressing WT, S224A or S241A LAT and normalized to mCherry expressing JLAT cells. Protein pulldown was further normalized to relative LAT pulldown with log-fold differences shown between groups. n=5 replicates/transductant.

Although we were unable to detect PLC*γ* and SLP-76 in our mass spectrometry screen, we confirmed a decrease in GADS binding to the S241A mutant, as observed in our co-immunoprecipitation experiment. We also observed a significant decrease in binding of the guanine nucleotide exchange factor Son of Sevenless 1 (Sos1), to S241A. Sos1 promotes LAT signalosome formation and is critical for T cell function.^29^ Interestingly, while some proteins showed decreased binding to the mutants compared to WT, a substantial number of proteins exhibited increased binding to the S224A and S241A mutants. This unexpected increase suggests that phosphorylation of these serines may not only regulate the binding of proteins to LAT but also influence the release of proteins. The exact mechanisms by which phosphorylation at these serines alters protein binding are still under investigation. Nonetheless, these data clearly confirm phosphorylation of S224 and S241 is a critical step in T cell signaling and may regulate T cell response to antigen.

### Phosphorylation at S224 and S241 is not required for LAT localization to the IS

We investigated whether phosphorylation of LAT serines S224 and S241 is required for LAT translocation and retention at the IS. **Figure S5C** displays LSCM images showing normal LAT translocation to the plasma membrane in J.LAT cells transduced with WT LAT following activation with αCD3/CD28. LAT mutants S224A, S241A, and the double mutant S2A also localized to the membrane similarly to WT LAT. This indicates that, although DNA-PKcs is essential for proper LAT localization to the IS, as demonstrated by NU7441 treatment and shRNA knockdown of DNA-PKcs, the phosphorylation of these two serines is not critical. These findings suggest that additional mechanisms mediate DNA-PKcs’s impact on LAT translocation.

Upon TCR activation, current studies suggest that LAT, located in plasma membrane lipid rafts and intracellular vesicles, traffics through the Golgi apparatus before being transported to the IS within secretory vesicles.^3,4,30^ Our observation that inhibiting DNA-PKcs prevents LAT accumulation at the IS, combined with previous findings that DNA-PKcs is essential for proper Golgi vesicle development, suggests DNA-PKcs may play a unique role in regulating Golgi-mediated trafficking.^31^ Therefore, we hypothesized that DNA-PKcs inhibition disrupts the trafficking of LAT-containing vesicles from the trans-Golgi to the IS. To investigate this, we analyzed the localization of DNA-PKcs within the Golgi compartments of T cells. Using a Golgi fractionation kit, we determined that active pDNA-PKcs localizes to both the cis and trans-Golgi compartments in activated T cells. Notably, pDNA-PKcs was found in secretory vesicles alongside LAT (**Figure S6A**). To support the hypothesis that DNA-PKcs inhibition affects Golgi-mediated trafficking of LAT, we compared LAT localization patterns in Jurkat T cells treated with Brefeldin A, a vesicle transport inhibitor.^32^ Brefeldin A treatment similarly attenuated LAT localization to the plasma membrane as observed with DNA-PKcs inhibitor treatment (**Figure S6B**). Other TCR-related proteins, such as CD3ζ and Lck, also traffic through the Golgi to localize to the IS.^33,34^ If DNA-PKcs influences this process, we would expect defects in the IS localization of these proteins as well. **Figure S6C** shows LSCM images of CD3ζ and Lck (p-Y394) in activated T cells. Consistent with LAT, the signal intensities of CD3ζ and p-Lck at the plasma membrane were also attenuated upon DNA-PKcs inhibition.

While the role of DNA-PKcs in the cis and trans-Golgi of resting and activated T cells remains largely unexplored, our data indicate a potential mechanism by which DNA-PKcs regulates the recruitment of proteins to the IS through the control of Golgi-mediated vesicular trafficking following antigen engagement.

### Generation of inducible T cell-specific DNA-PKcs deleted mice

To thoroughly understand the role of DNA-PKcs in T cells, it is imperative to investigate DNA-PKcs function in mature primary T cells. However, animals with inactivating mutations in DNA-PKcs during embryonic development have a significant reduction in mature B and T cells. This phenotype hinders investigations into the function of DNA-PKcs in mature T cells. To analyze its activity specifically in mature T cells, we generated two unique mouse models in which DNA-PKcs is genetically deleted specifically in either CD4⁺ or CD8⁺ T cells.

Floxed DNA-PKcs (DNA-PKcs^fl/fl^) mice, obtained from Dr. Jay H. Chung, were bred with either the tamoxifen-inducible CD4-Cre mouse strain to generate Cre^+^CD4-PKcs^fl/fl^ mice, or with the tamoxifen-inducible E8ICreERT2-CD8 mouse strain, received from Dr. Dario Vignali, to generate Cre^+^CD8-PKcs^fl/fl^ mice (**Figure S7A**).^35,36^ Control mice in this study expressed Cre recombinase but WT DNA-PKcs (Cre^+^CD4-PKcs^wt/wt^, Cre^+^CD8-PKcs^wt/wt^). Eight-week-old Cre^+^CD4-PKcs^wt/wt^, Cre^+^CD4-PKcs^fl/fl^, Cre^+^CD8-PKcs^wt/wt^ or Cre^+^CD8-PKcs^fl/fl^ mice with a mature immune system were treated with tamoxifen for seven consecutive days. DNA-PKcs expression was then analyzed by qPCR and western blot in CD4⁺ or CD8⁺ T cells isolated from spleens. Tamoxifen treatment resulted in a significant reduction of DNA-PKcs expression (up to 90%), which persisted for a minimum of 30 days (**Figure S7B**). Loss of DNA-PKcs expression within CD4⁺ or CD8⁺ T cells did not alter mouse activity or affect splenic size or weight (**Figure S7C**). Loss of DNA-PKcs expression did not impact baseline levels of naïve CD4⁺ and naïve CD8⁺ T cells, B cells, macrophages or dendritic cells isolated from the spleens of tamoxifen-treated mice (**Figure S7D**).

### DNA-PKcs-deficient CD4^+^ and CD8^+^ T cells have functional deficits

#### Reduced Cytokine Production

To evaluate how the loss of DNA-PKcs expression affects T cell function, we first assessed the ability of DNA-PKcs-deficient T cells to produce cytokines in response to antigen presentation using mixed lymphocyte reactions (MLRs). Splenocytes isolated from Balb/c mice and treated with mitomycin C served as stimulators, while control or DNA-PKcs-deficient CD4⁺ or CD8⁺ T cells isolated from our transgenic mice acted as responders. Loss of DNA-PKcs expression significantly reduced the production of IL2 and IFN*γ* by CD8⁺ T cells following stimulation with allogeneic antigens (**Figure 5A**). Reduced cytokine expression was also observed in DNA-PKcs-deficient CD8^+^ T cells activated with αCD3/CD28 (**Figure 5A**). Loss of DNA-PKcs expression in CD4^+^ T cells did not affect IL2 production by either alloantigen or αCD3/CD28 stimulation. However, IFN*γ* was increased in DNA-PKcs-deficient CD4⁺ activated with αCD3/CD28 but not alloantigens (**Figure 5A**). Effects on cytokine production were not attributable to impaired T cell proliferation or increased apoptosis. Loss of DNA-PKcs expression did not affect the proliferation or enhance apoptosis in either αCD3/CD28-activated CD4⁺ or CD8⁺ T cells, as assessed by CellTrace proliferation assays and Annexin V staining (**Figure S7F**). This highlights that, although chemical inhibition of DNA-PKcs impairs T cell proliferation, the residual levels of DNA-PKcs present in our transgenic T cells may be sufficient to support normal proliferation.^37^ Taken together, these findings suggest DNA-PKcs is critical for the function of CD8^+^ T cells but less so in CD4^+^ T cells.

**Figure 5.**
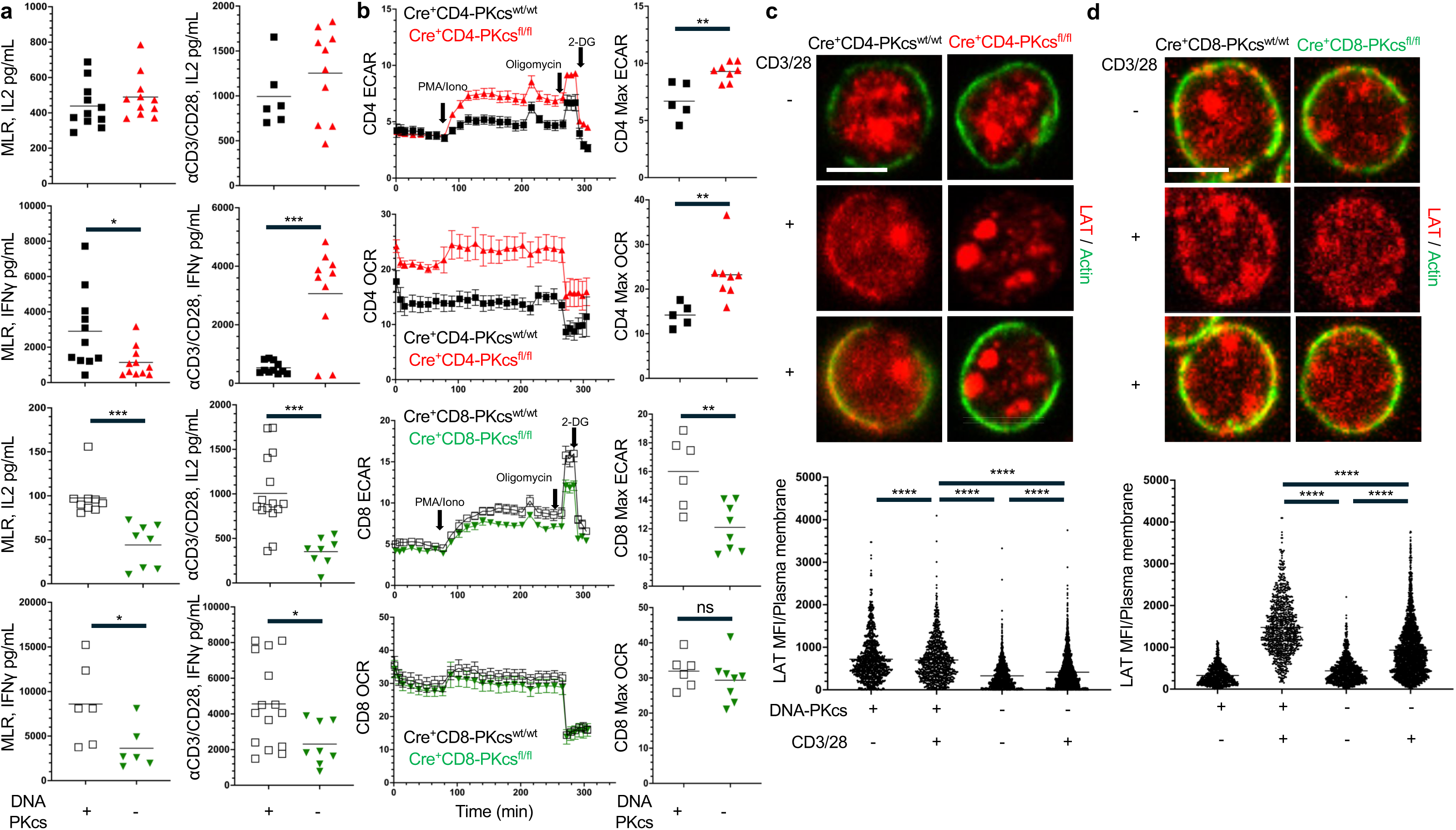
DNA-PKcs-deficient CD4^+^ and CD8^+^ T cells have functional deficits and impaired LAT localization at the plasma membrane. **A)** Isolated CD8^+^ T cells from Cre⁺CD8-PKcs^wt/wt^ and Cre^+^CD8-PKcs^fl/fl^ mice and isolated CD4^+^ T cells from Cre⁺CD4-PKcs^wt/wt^ and Cre^+^CD4-PKcs^fl/fl^ mice were stimulated for 48 hours by either αCD3/CD28 or by coculture with splenocytes from BALB/c mice (MLR, 4:1 donor to responder). Supernatant was analyzed for IL2 and IFNγ by ELISA. **B)** Representative ECAR and OCR of isolated CD4^+^ and CD8^+^ T cells. Basal metabolic rates were recorded for 80 minutes, followed by sequential stimulation with PMA and Ionomycin to activate T cells. Metabolic inhibitors, oligomycin (ATP-synthase inhibitor) and 2-DG, glycolysis inhibitor, were used as internal controls. Maximal ECAR and OCR values were determined post-stimulation. LSCM of isolated **C)** CD4^+^ T cells and **D)** CD8^+^ T cells stimulated for 2-minutes by αCD3/CD28 with quantification of LAT (red) signal at the plasma membrane. Scale bar = 5 µm. Representative images from n=3 independent experiments. **A,B)** unpaired T-test and **C,D)** One factor ANOVA + Tukey’s multiple comparisons used to determine statistical significance (α=0.05, *p<0.05, **p< 0.01, ***p< 0.0005, ****p< 0.0001), error bars represent SEM, n= at least 6 mice/group.

#### Impaired Metabolic Reprogramming

T cell activation induces metabolic reprogramming characterized by a switch from oxidative phosphorylation to glycolysis. Therefore, we analyzed changes in extracellular acidification rate (ECAR), an indicator of aerobic glycolysis, in our DNA-PKcs-deficient primary T cells.^38^ Basal respiration rates were similar between WT and DNA-PKcs-deficient CD8⁺ T cells; however, upon stimulation with PMA/Ionomycin, DNA-PKcs-deficient CD8⁺ T cells exhibited reduced ECAR and OCR (oxygen consumption rate) rates compared to WT counterparts, indicating impaired metabolic reprogramming. (**Figure 5B**). Interestingly, loss of DNA-PKcs expression in CD4⁺ T cells resulted in a significant increase in ECAR and OCR compared to WT upon PMA/Ionomycin stimulation (**Figure 5B**). While this effect differs from that observed in CD8⁺ T cells, the higher ECAR in CD4⁺ T cells could indicate a compensatory upregulation of glycolytic pathways or an alternative metabolic adaptation in response to DNA-PKcs deficiency. While research is needed to understand the mechanisms behind this metabolic shift and its impact on CD4⁺ T cell function, these observations further validate a critical role for DNA-PKcs in T cells.

### LAT Localization to the Plasma Membrane is Reduced in Activated DNA-PKcs-Deficient Primary CD4^⁺^ and CD8^⁺^ T Cells

To further confirm the role of DNA-PKcs in LAT localization to the plasma membrane following T cell activation, as observed in Jurkat T cells, we analyzed LAT expression patterns in splenic CD4⁺ and CD8⁺ T cells isolated from Cre^+^CD4-PKcs^fl/fl^ and Cre^+^CD8-PKcs^fl/fl^ mice using LSCM. As expected, αCD3/CD28 activation in control T cells induced LAT expression at the plasma membrane of mouse T cells. However, LAT expression at the plasma membrane was significantly reduced in both DNA-PKcs-deficient CD4⁺ and CD8⁺ T cells (**Figures 5C, D**). This indicates that DNA-PKcs plays a crucial role in LAT trafficking to the plasma membrane following TCR stimulation.

Next, we examined how the loss of DNA-PKcs expression affects LAT tyrosine phosphorylation. In Jurkat T cells, we determined that phosphorylation of LAT at S241 is required for phosphorylation of tyrosines Y132, Y171, and Y191. However, inhibition of DNA-PKcs with the specific inhibitor NU7441 did not impact tyrosine phosphorylation, suggesting that S241 is likely phosphorylated by other kinases. Consistent with this finding, loss of DNA-PKcs expression in CD4⁺ or CD8⁺ T cells did not affect phosphorylation at Y132 and Y191 (**Figures S8A, B**). These results suggest a dual role for DNA-PKcs in T cells: it is critical for the trafficking of proteins to the IS and influences signaling from the LAT signalosome.

### Loss of DNA-PKcs Results in Dysfunctional T Cells Unable to Control Tumor Growth

T cells are a central component of eliminating cancerous cells, therefore, to confirm DNA-PKcs as a pivotal player in T cell activity and their ability to mount an effective immune response against tumors, we analyzed the effect of DNA-PKcs-deficient CD4⁺ or CD8⁺ T cells on tumor growth *in vivo*. Following tamoxifen treatment and injection of MC38 colorectal cancer cells, Cre^+^CD8-PKcs^fl/fl^ mice had significantly accelerated tumor growth resulting in reduced survival compared to Cre⁺CD8-PKcs^wt/wt^ (control) mice with DNA-PKcs-expressing CD8⁺ T cells (**Figures 6A, B**) This finding was further validated with B16F10 melanoma tumor, where survival decreased and tumor growth was accelerated in Cre^+^CD8-PKcs^fl/fl^ mice compared to controls (**Figures 6C, D**). These results demonstrate that loss of DNA-PKcs in CD8⁺ T cells leads to dysfunctional T cells unable to effectively control tumor growth. Interestingly, MC38 tumors grew at an accelerated pace during initial response in Cre^+^CD4-PKcs^fl/fl^ mice where T cell control is most evident, however tumor growth eventually equalized to control mice, resulting in no significant difference in survival (**Figure 6E, F**). This further suggests DNA-PKcs plays a different regulatory role in CD4^+^ versus CD8^+^ T cells.

**Figure 6.**
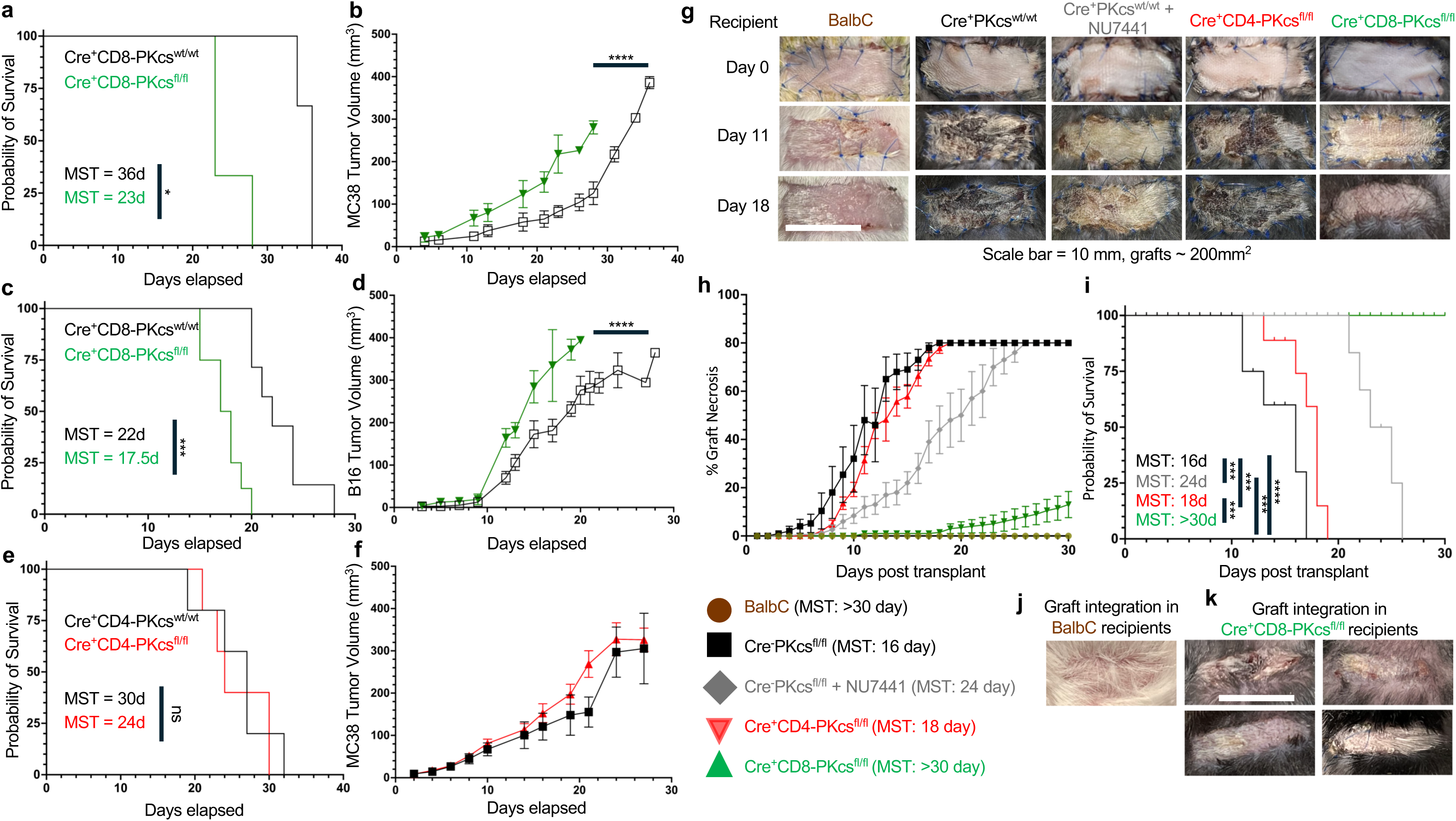
DNA-PKcs-deficient CD8^+^ T cells cannot control tumor growth or respond to allografts. **A,B)** MC38 adenocarcinoma cells or **C,D)** B16 melanoma cells were injected into rear flank of Cre⁺CD8-PKcs^wt/wt^ and Cre⁺CD8-PKcs^fl/fl^ mice. **A)** MC38 and **C)** B16 tumor cells in Cre^+^CD8-PKcs^fl/fl^ mice had accelerated growth compared to Cre⁺CD8-PKcs^wt/wt^ mice (****p<0.0001, nonlinear regression, error bars represent SEM, n=3 and 8/group respectively). **B)** Cre^+^CD8-PKcs^fl/fl^ mice exhibited decreased median survival time (MST: 23d) compared to Cre⁺CD8-PKcs^wt/wt^ mice (MST: 36d) (*p<0.05, log-rank analysis). **D)** Cre^+^CD8-PKcs^fl/fl^ mice exhibited decreased MST (17.5d) compared to Cre⁺CD8-PKcs^wt/wt^ (22d) (***p<0.0005, log-rank analysis). MC38 adenocarcinoma cells were injected into the rear flank of Cre⁺CD4-PKcs^wt/wt^ and Cre^+^CD4-PKcs^fl/fl^ mice. **E)** Tumor volume in Cre^+^CD4-PKcs^fl/fl^ mice had a similar rate of growth compared to Cre⁺CD4-PKcs^wt/wt^ mice (*p<0.05, nonlinear regression, error bars represent SEM, n=6/group). **F)** Cre^+^CD4-PKcs^fl/fl^ mice exhibited similar MST (24d) compared to Cre⁺CD4-PKcs^wt/wt^ mice (27d) (p=0.74, log-rank analysis). BALB/c tail skin allografts were transplanted onto backs of syngeneic BALB/c (n=5), or fully allogeneic Cre⁺PKcs^wt/wt^ (n=5), Cre⁺-PKcs^wt/wt^ + NU7441 (n=4), Cre^+^CD4-PKcs^fl/fl^ (n=7), or Cre^+^CD8-PKcs^fl/fl^ (n=10) recipient mice. **G)** Representative examples of skin graft healing on day 0 (day of surgery), 11 days post-op and 18 days post-op. **H)** Graft necrosis was measured daily with DNA-PKcs inhibitor NU7441 slowing graft necrosis progression. Cre^+^CD8-PKcs^fl/fl^ recipient mice had no necrosis until integration where minor necrosis (<20%) was apparent in 2 of 10 mice, 8 of 10 mice did not show any necrosis. **I)** Compared to control mice (MST: 16d), median survival of allograft was significantly increased in Cre⁺-PKcs^wt/wt^ in NU7441 treated mice (MST: 24d, p<0.0001) and Cre^+^CD8-PKcs^fl/fl^ recipient mice (MST: >30d, p<0.0001) and marginally increased in CD4-PKcs^-/-^ mice (MST: 18d, p<0.05), compared by log-rank analysis.**J)** Syngeneic BALB/c and **K)** Cre^+^CD8-PKcs^fl/fl^ recipient mice had successful graft integration and graft hair growth by day 30.

### DNA-PKcs-deficient CD8^⁺^ T Cells do not reject allogeneic skin grafts

Cytotoxic T cells are the primary drivers of graft rejection in transplantation. We recently published that chemical inhibition of DNA-PKcs reduced T cell-mediated rejection and extended graft survival in a murine allogeneic skin graft model.^39^ We used this model to analyze defects in DNA-PKcs-deficient T cell response to alloantigen. **Figure 6G** contains representative images of skin grafts. Grafts were considered completely rejected, as seen in the graft from control mice (PKcs^wt/wt^) on Day 18, when necrosis consumed 80% of the entire graft. Treatment with the DNA-PKcs inhibitor NU7441 significantly extended graft survival compared to vehicle treated controls, supporting our previous findings (**Figure 6H**). Although loss of DNA-PKcs in CD4⁺ T cells did not significantly alter the overall rejection timeframe, there was an initial delay in rejection that accounts for a minimal increase in survival. For instance, average graft necrosis in control mice was 30% by Day 8 but less than 10% in Cre^+^CD4-PKcs^fl/fl^. This trend continued until around Day 15 when necrosis rates were similar between control and Cre^+^CD4-PKcs^fl/fl^. However, loss of DNA-PKcs in CD8⁺ T cells extensively attenuated graft necrosis and in most cases completely prevented any rejection (**Figure 6I**). Graft health was preserved for over 30 days with only sparse necrosis observed in Cre^+^CD8-PKcs^fl/fl^ recipients. Furthermore, over 50% of Cre^+^CD8-PKcs^fl/fl^ recipients had complete graft integration with hair growth by day 25 (**Figures 6J, K**). This exciting finding supports the tumor growth data and indicates that DNA-PKcs is required for the cytotoxic response to alloantigens.

## Discussion

Our study elucidates the multifaceted role of DNA-PKcs in T cell biology, extending beyond its well-established function in DNA repair. We have uncovered that DNA-PKcs is integral for both trafficking and signaling mechanisms within mature T cells, specifically influencing the localization and function of the immune synapse-associated protein LAT.

We show that DNA-PKcs is localized to the IS following TCR stimulation. This intriguing finding suggests DNA-PKcs has a critical function in regulating T cell receptor signal transduction induced upon antigen recognition. This corresponds with previous reports indicating that DNA-PKcs localized to lipid rafts within the plasma membrane of tumor cells promotes intracellular receptor signal transduction initiated by ionizing radiation.^40^ Similar to our findings, these results were unique to DNA-PKcs and not seen with other DNA damage repair kinases. This study, along with our data, supports a function for DNA-PKcs in receptor-mediated signal transduction, specifically TCR signaling in T cells. The mechanism by which DNA-PKcs is activated and recruited to the IS following stimulation is unknown. We are actively investigating these processes including the involvement of the KU70/80 complex in the activation and trafficking of DNA-PKcs to the IS.

One pivotal finding of our research is the identification of DNA-PKcs as a critical regulator of LAT trafficking to the IS. Upon TCR activation, DNA-PKcs is activated and localizes to the IS. Activation of DNA-PKcs is essential for the translocation of LAT to the IS, a process that is impaired upon inhibition or genetic deletion of DNA-PKcs. Interestingly, while phosphorylation of LAT S224 and S241 is crucial for IL2 production and protein binding, it does not appear to be necessary for LAT’s localization to the plasma membrane. Our Golgi fractionation and Brefeldin A experiments support a hypothesis that DNA-PKcs regulates Golgi-mediated vesicular transport of LAT, and IS associated proteins, from the trans Golgi to the IS.

We show that phosphorylation of LAT at S224 and S241 is essential for the phosphorylation of LAT Y132, Y171, and Y191 which facilitates assembly of the LAT signalosome. S224 and S241 reside in the unstructured C-terminus tail of LAT distal to transmembrane domain and helix structure. While tyrosines have been the primary phosphorylation sites studied within LAT, other serine phosphorylation sites have been identified within LAT that are required for protein binding and proximal signaling.^41^ Our mass spectrometry results suggests that phosphorylation of S241 regulates which proteins bind to LAT as well as the release of proteins during T cell activation. This was evident by the increased binding of several proteins to S241 compared to WT LAT. This is critical given recent findings from the Weiss lab showing temporal LAT protein binding effects antigen discrimination in T cells.^42^ Future studies should focus on determining the role of LAT serine phosphorylation specifically S241 in TCR specificity, tolerance to self-antigens, and T cell response outcome.

Our *in vivo* studies using Cre^+^CD4-PKcs^fl/fl^ and Cre^+^CD8-PKcs^fl/fl^ mouse models reveal a differential impact of DNA-PKcs deficiency on CD4⁺ versus CD8⁺ T cells although similar deletion of DNA-PKcs was observed in both models. While CD8⁺ T cells lacking DNA-PKcs displayed significant defects in controlling tumor growth and rejecting skin grafts, DNA-PKcs-deficient CD4⁺ T cells did not exhibit similar levels of dysfunction although they did display alterations in LAT IS localization. This disparity suggests that DNA-PKcs plays a more critical role in the cytotoxic functions of CD8⁺ T cells compared to the helper functions of CD4⁺ T cells. Previous studies showed reduced expression of DNA-PKcs prevents Th1 or Th2 CD4^+^ T cell differentiation suggesting additional evaluation of the CD4 model is needed to determine the effects of DNA-PKcs on the CD4^+^ T cell population.^19^ It should be noted that LAT is more highly expressed in human T cells than mouse T cells and could have more of an impact on human T cell function.^43^ With our transgenic model fully developed and validated, we can continue to explore the function of DNA-PKcs in T cell signaling potentially identifying changes in T cell subpopulations that account for the attenuated response to antigen.

DNA-PKcs inhibitors have been widely studied for the ability to sensitize tumor cells to chemotherapy and radiation therapy.^44,45^ The impaired signaling in DNA-PKcs-deficient T cells translate to diminished effector functions such as cytokine production and cytotoxic activity, which are crucial for anti-tumor immunity. These findings indicate DNA-PKcs inhibitors are a double edge sword for cancer therapy: inhibition weakens tumor cells but also diminishes T cell immunity. This accounts for the mild benefits of DNA-PKcs inhibitors in cancer clinical trials where the best response is stable disease despite significant results in rodent models.^46–48^ Alternatively, this study supports our previous publication highlighting the potential benefit of DNA-PKcs inhibitors as anti-rejection therapy for transplant patients where loss of T cell immunity improves graft survival.^49^

In summary, our findings position DNA-PKcs as a pivotal regulator of T cell function, bridging the processes of protein trafficking and signalosome assembly. By modulating both the localization of key signaling proteins and the downstream signaling pathways, DNA-PKcs orchestrates effective T cell responses essential for immune surveillance and pathogen defense. These insights not only advance our understanding of T cell biology but also open new avenues for therapeutic interventions targeting DNA-PKcs in immune-related diseases.

## Supplementary figure legends

**Supplementary Figure 1. T cell receptor (TCR) stimulation activates DNA-PKcs. A)** Continuous αCD3/CD28 TCR stimulation of E6.1 Jurkat T cells with αCD3/CD28 induces autophosphorylation of DNA-PKcs. **B)** Temporary (2 min) TCR stimulation of E6.1 Jurkat T cells induces time sensitive elevation in DNA-PKcs phosphorylation.

**Supplementary Figure 2. DNA-PKcs phosphorylates LAT at serine 224 and 241.** *In vitro* kinase assay analysis of DNA-PKcs enzyme phosphorylation of the **A)** control peptide or **B)** LAT peptide. **C**) Mass spectrometry analysis of LAT peptide following *in vitro* kinase assay with serine sites on LAT +/-DNA-PKcs. **D)** Sequence alignment of LAT from mice to humans showing conservation of serine residues. Error bars=SEM. One factor ANOVA + Tukey’s multiple comparisons used for statistical testing. (α=0.05, *p<0.05, **p< 0.01, ***p< 0.0005, ****p< 0.0001).

**Supplementary Figure 3. DNA-PKcs inhibition diminishes LAT localization at the plasma membrane: A)** 2-minute TCR stimulation of isolated human T cells, from whole blood PBMC’s. LSCM shows LAT (red) localization at the plasma membrane with pDNA-PKcs (green) upon TCR stimulation which is attenuated with NU7441 (5 µM). **B)** Quantification of LAT expression shows significant increase in mean fluorescent intensity (MFI) upon TCR stimulation with significant decrease when DNA-PKcs is inhibited. Scale bar = 5 µm. Representative images from n=3 independent experiments. One factor ANOVA + Tukey’s multiple comparisons used (α=0.05, ****p<0.0001).

**Supplementary Figure 4. Validation of J.LAT transductants.** All vectors include mCherry fluorescent protein either independently (JLAT + mCherry) or tagged via flexible linker to LAT (JLAT + WT LAT, S224A LAT, S241A LAT and S2A LAT). **A)** mCherry expression was validated by flow cytometry with each mCherry transductant showing detectable expression over baseline**. B)** LAT protein expression was validated by western blot.

**Supplementary Figure 5. Phosphorylation of S224 and S241 are required for IL2 production but not proliferation or LAT translocation. A)** JLAT transductants show no difference in proliferative capacity measured by Cell Trace Violet staining. **B)** Generations were produced at similar rates with no difference between cell lines (simple linear regression, p=0.46). **C)** LAT serine phosphorylation sites are not required for LAT localization to the plasma membrane as **D)** greater than 90% of LAT signal is at the plasma membrane across transductants with 2-minute αCD3/CD28 stimulation. Scale bar = 5 µm. Representative images from n=3 independent experiments. One factor ANOVA + Tukey’s multiple comparisons used to determine statistical significance (α=0.05).

**Supplementary Figure 6. DNA-PKcs localizes with LAT in Golgi secretory vesicles in T cells.** E6.1 Jurkat T cells were stimulated with αCD3/CD28 for 15 minutes and then lysed with Golgi fractionation buffer. **A)** Total and phosphorylated DNA-PKcs (pDNA-PKcs) are present in both the cis- and trans-Golgi fractions. Upon TCR stimulation, the secretory fraction exhibits an increase in pDNA-PKcs expression, in the presence of LAT. **B)** LAT expression at the plasma membrane is attenuated by both DNA-PKcs inhibition (NU7441) and inhibition of secretory vesicle blebbing (Brefeldin). **C)** Inhibition of DNA-PKcs prevents early TCR signaling markers like pLck and CD3ζ who undergo similar trafficking to the plasma membrane, from localizing to the plasma membrane. Scale bar = 5 µm. Representative images from n=3 independent experiments. One factor ANOVA + Tukey’s multiple comparisons (α=0.05, ****p<0.0001).

**Supplementary Figure 7. Generation and validation of inducible T cell specific DNA-PKcs deleted mice. A)** Schematic of breeding strategy producing Cre⁺CD4-PKcs^wt/wt^, Cre^+^CD4-PKcs^fl/fl^, Cre⁺CD8-PKcs^wt/wt^ and Cre^+^CD8-PKcs^fl/fl^ mice. Following 7-day tamoxifen treatment via oral gavage at 10mg/kg, **B)** DNA-PKcs mRNA and protein is significantly decreased in isolated CD4^+^ and CD8^+^ T cells. **C)** Spleen size and weight do not differ between control and DNA-PKcs knockout mice (spleen weight normalized to body weight**). D)** Spleens from control and DNA-PKcs-deficient mice contain similar numbers of total splenocytes and CD4^+^ or CD8^+^ isolated T cells following red blood cell lysis. **E)** Total splenocyte populations including CD4^+^ T cells, CD8^+^ T cells, CD19^+^ B cells, F4/80^+^ macrophages and CD11c^+^ dendritic cells showed no difference in levels across Cre⁺CD8-PKcs^wt/wt^, Cre^+^CD8-PKcs^fl/fl^, Cre⁺CD4-PKcs^wt/wt^ and Cre^+^CD4-PKcs^fl/fl^ mice. **F)** CD4^+^ and CD8^+^ T cells have similar levels of apoptotic cell death within 48 hours of αCD3/CD28 TCR stimulation. **G)** There is no difference in proliferative capacity of CD4^+^ or CD8^+^ T cells between control and DNA-PKcs-deficient mice as shown by similar population generation with Cell Trace violet stain. Representative data from at least n=6 mice/treatment group. Unpaired T-tests used to determine statistical significance (α= 0.05) between control and KO cells for DNA-PK knockdown, spleen weight comparison, and T-cell subset comparison, apoptotic cell death measured with paired T-test for 0 hr and 48 hr repeated measure between PKcs^wt/wt^ and PKcs^fl/fl^ mice.

**Supplementary Figure 8. DNA-PKcs-deficient CD4^+^ and CD8^+^ T cells have normal LAT tyrosine phosphorylation levels**. Cre⁺CD4-PKcs^wt/wt^, Cre^+^CD4-PKcs^fl/fl^, Cre⁺CD8-PKcs^wt/wt^ and Cre^+^CD8-PKcs^fl/fl^ mice were tamoxifen treated for 7 days prior to CD4^+^ or CD8^+^ T cell splenocyte isolation. **A)** CD4^+^ T cells and **B)** CD8^+^ T cells were stimulated with αCD3/CD28 for two minutes and phosphorylation of tyrosine residues were investigated via immunoblot. DNA-PKcs deficiency alone did not prevent tyrosine phosphorylation in CD4^+^ or CD8^+^ T cells. Percentage of TCR stimulated LAT phosphorylation was normalized to resting pLAT samples via imageJ analysis. n=3 mice per group.

## Resource availability

### Lead contact

Requests for further information and resources should be directed to and will be fulfilled by the lead contact Dr. Marie Burdine

### Materials availability

Reagents and resources described in this paper will be available from the lead contact upon reasonable request and after completion of a Materials Transfer Agreement.

### Data and code availability

The mass spectrometry proteomics data have been deposited to the ProteomeXchange Consortium via the PRIDE partner repository.

This paper does not report original code.

Any additional information required to reanalyze the data reported in this paper is available from the lead contact upon request.

## Acknowledgements

This work was performed in collaboration with the UAMS IDeA National Resource for Quantitative Proteomics (NIH R24GM137786). Authors would like to acknowledge Dr. Jay H. Chung at NHLBI for the DNA-PKcs floxed mice, Dr. Dario Vignali at University of Pittsburgh for the E8ICreERT2-CD8 mouse strain, and the Arthur Weiss laboratory at UCSF for the J. LAT cell line.

This work was supported by NIH 1R21AI173759-01A1 (MSB, BK), Arkansas Children’s Research Institute (MSB) and the Division of Surgical Research (MSB). Additional funding was provided by NIH/NIGMS T32-GM106999 (RRR). The content is solely the responsibility of the authors and does not necessarily represent the official views of the NIH.

## Author Contributions

MSB and LB conceived the study and obtained funding. RRR, ACA, LB and MSB designed all of the experiments. RRR and ACA performed and interpreted all of the experiments with assistance from MSB and LB. ZJW performed *in vitro* kinase assay and mass spectrometry study on LAT peptide. NAC performed skin transplant studies. BHH maintained all animal strains, performed genotyping and analysis of impact of DNA-PKcs deletion on immune cells. BK assisted with metabolism studies. AJS assisted with the analysis of mass spectrometry data and provided statistical oversight. RRR, ACA, ZJW, NAC, BH, BK, AJS, LB and MSB analyzed and interpreted the data. MSB and RRR wrote the manuscript with input from all of the authors.

## Methods

### Cell Culture, antibodies and reagents

Jurkat E6.1 human T cell leukemia and Raji B cells were obtained from American Type Culture Collection (ATCC). LAT deficient Jurkat T cells, J. LATs, were kindly provided by the Arthur Weiss, MD, PhD (UCSF). Cells were maintained in RPMI 1640 with 10% FBS and 1% Penicillin and streptomycin (P/S). Tumor cells lines B16-F10 and MC-38 were provided by Brian Koss, PhD (UAMS). B16-F10 cells were maintained in DMEM with 10% FBS and 1% P/S. MC-38 cells were cultured with 10% FBS, 1% P/S, 0.1mM non-essential amino acids, 1mM sodium pyruvate and 10mM HePES. Mouse primary T cells were maintained in RPMI 1640 with 1x NEAA, 1x sodium pyruvate and 55 µM BME. Human PBMCs were ordered from Stem cell Technologies and maintained in RPMI 1640 with 10% FBS and 1% Penicillin and streptomycin. See complete list of antibodies, shRNA’s, and inhibitors in supplementary material.

### Western blot, cell fractionation and co-immunoprecipitation

Cells were harvested with ice-cold PBS and placed on ice. Cells were spun down at 500xg for 10 minutes, washed with ice cold PBS (phosphate-buffered saline) prior to lysis. Whole cell lysate was collected using RIPA lysis buffer. For cytosolic extraction, cells were lysed with 0.1% NP-40 (+ protease and phosphatase inhibitors) on ice for 10 minutes. Cells were then centrifuged at 10,000xg for 20 minutes to remove nuclear compartment. Supernatant was then used for western blot analysis. Co-immunoprecipitation was completed using cytosolic extraction followed by primary antibody incubation and protein A/G bead pulldown. Input sample was collected from the cytosolic extract.

### Laser scanning confocal microscopy (LSCM)

Micro-slide 8-well glass-bottom coverslips (Ibidi, Cat: 80807) were coated with 0.01% poly-L-lysine (Poly-Lys, Sigma-Aldrich, P8920) in PBS ± CD3 for 2 hours at 37°C. T cells were seeded at a concentration of 250,000 cells/ml onto the coated coverslips with or without soluble CD28 for the specified duration. After incubation, cells were fixed with 4% paraformaldehyde for 10 minutes, followed by a single wash with PBS. The cells were then permeabilized using 0.1% Triton X-100 in PBS for 20 minutes and washed once more with PBS. To block nonspecific binding, cells were incubated in 2.5% BSA in TBST for 1 hour. Cells were incubated with primary and secondary antibodies of interest, washed three times with TBST and stained with Hoechst. Imaging was performed using a Zeiss LSM scanning confocal microscope. Analysis of images was completed using the ZenBlue software.

### Immune conjugate formation

Raji B cells were washed, resuspended at a concentration of 1 × 10^6^ cells/mL in RPMI without FCS and labeled with Hoechst dye (Thermo Fisher, C2110) for 20 min at 37 °C. Labeling was stopped with RPMI 10% FCS and cells were washed once and resuspended at 1 × 10^6^ cells/mL. Cells were pulsed with Recombinant Staphyloccocus Enterotoxin Type E (SEE, Cellgenetech, MBS1112600) (1 µg/mL) or left untreated for 30 min at 37°C before being washed once and resuspended at a concentration of 1 × 10^6^ cells/mL. 100,000 Raji cells were incubated on coverslips for 30 min, washed once with warmed PBS and 150,000 Jurkat cells were added for 30 min. Cells were fixed, permeabilized and blocked similar to aforementioned imaging.

### Cellfree kinase assay

The DNA-PK Kinase Enzyme System (Cat # V4106) and ADP-Glo Assay (Cat # V6930) from Promega were used as previously described (Waldrip et al., 2021). Briefly, DNA-PK enzyme was incubated with control peptide or LAT peptide in a 96-well plate format with 150 μM ATP, 1 μg of peptide. DNA-PK enzyme was used at 50 units per reaction, in a reaction volume of 25 μl. Enzyme, peptide, and ATP were incubated for 1 hour at room temperature prior to ADP-Glo protocol. Luminescence was measured on synergy HTX multiplate reader (BioTek) using an integration time of 0.25 second per well. Enzyme plus ATP samples, no peptide, were used to determine baseline luminescence since DNA-PKcs is known to phosphorylate itself.

### Development of stable LAT phosphomutant cell lines

The CRISPR knockout LAT cell line, J.LAT, was developed, validated and kindly supplied by the Arthur Weiss laboratory (UCSF). Transfer plasmids containing mCherry fluorescent probe and mCherry tagged WT LAT, S224A LAT, S241A LAT or S224/241A LAT were developed, validated and purchased from vectorbuilder. Envelope and packing plasmids from Addgene were transfected into HEK293T cells with transfer plasmids to produce lentiviral particles which were then used to transduce J.LAT cells. J.LAT transductants were screened by their mCherry expression by flow cytometry and cells were grown in the presence of puromycin (2 µg/mL). Transduction of LAT version was validated by PCR (plasmidosaurus). LAT expression was validated across mutants by western blot.

Non-replicative VSV-g pseudotyped lentiviral particles were produced by transfecting HEK-293T cells with Gag, Pol, rev, encoding plasmid (pPAX2), envelop encoding plasmid (pMD2.G) and either the WT LAT or Mutant LAT construct. Mutant and WT LAT lentiviral transfer plasmids were validated through vectorbuilder quality control prior to use *in vitro*. Mutant and WT LAT were expressed using the pcDNA3 lentiviral expression system which also encodes a puromycin-resistance gene for production of stable lines. LAT transfer plasmids were combined with packaging and envelope plasmids and transfected into HEK293T cells in antibiotic free DMEM (10% FBS) according to published methods. 48 hours later, viral supernatant was collected from HEK293T cell culture and passed through a 0.2 µM filter. Viral media was incubated the with J.LAT cells via spinfection (2000 rpm for 1 hr) with polybrene (5 µg/mL) and incubated at 37°C for 24 hours. Viral media was removed and transduced cells were replenished with fresh RPMI and allow to grow for 2 days prior to puromycin addition (2 µg/mL). MCherry expression (tagged to the c-terminus of LAT) was measured weekly.

### Transfection

Jurkat T cells were transfected with shRNA to DNA-PKcs using Lonzo electroporation system. Following pulse, Jurkat T cells were plated in fresh, supplemented RPMI and incubated at 37°C for 36 hours prior to downstream analysis. Knockdown of protein expression was validated on western blot.

### Golgi analysis in T cells

E6.1 Jurkat T cells were cultured under normal conditions and plated at 1 x10^6^ cells/mL. Cells were harvested and washed with ice cold PBS and pellets were used in the Golgi apparatus enrichment kit (cat: GO-037, Invent Biotechnologies). Following initial lysis with kit lysis buffer, 15% of sample volume was collected for input. This kit separates cis Golgi from the secretory trans Golgi and with kit recommended controls of GM130 and CLINT1, respectively. Following protocol from manufacture, these fractions were collected and run on western blot. For LSCM analysis of the Golgi, cells were treated with Brefeldin (10 µg/mL) for 4 hours before activation.

### Co-Immunoprecipitation Mass Spectrometry

Following the co-immunoprecipitation protocol as outlined above, lysates were submitted to the UAMS proteomics core for analysis via the following:

### FASP Methods – Orbitrap Exploris DIA

Protein samples were reduced, alkylated, and digested on-bead using filter-aided sample preparation with sequencing grade modified porcine trypsin (Promega).^50^ Tryptic peptides were then separated by reverse phase XSelect CSH C18 2.5 um resin (Waters) on an in-line 150 x 0.075 mm column using an UltiMate 3000 RSLCnano system (Thermo). Peptides were eluted using a 60 min gradient from 98:2 to 65:35 buffer A:B ratio. Eluted peptides were ionized by electrospray (2.4 kV) through a heated capillary (275°C) followed by data collection on an Orbitrap Exploris 480 mass spectrometer (Thermo Scientific). Precursor spectra were acquired with a scan from 385-1015 Th at a resolution set to 60,000 with 100% AGC, max time of 50 msec, and an RF parameter at 40%. DIA was configured on the Orbitrap 480 to acquire 40 x 15 Th isolation windows, normalized AGC target 500%, maximum injection time 40 ms). A second DIA was acquired in a staggered window (15 Th) pattern with optimized window placements from 390-990 Th.

Buffer A = 0.1% formic acid, 0.5% acetonitrile

Buffer B = 0.1% formic acid, 99.9% acetonitrile

Following data acquisition, data were searched using Spectronaut (Biognosys version 19.1) against the UniProt *Homo sapien* database (May 2024) using the direct DIA method with an identification precursor and protein q-value cutoff of 1%, generate decoys set to true, the protein inference workflow set to maxLFQ, inference algorithm set to IDPicker, quantity level set to MS2, cross-run normalization set to false, and the protein grouping quantification set to median peptide and precursor quantity. Protein MS2 intensity values were assessed for quality using ProteiNorm.^51^ The data was normalized using Log_2_ Median Intensities and analyzed using proteoDA to perform statistical analysis using Linear Models for Microarray Data (limma) with empirical Bayes (eBayes) smoothing to the standard errors. Proteins with an FDR adjusted p-value < 0.05 and a fold change > 2 were considered significant. Heatmaps were constructed using R version 4.3.0 and the following packages: tidyverse, ggsci, and cowplot.

### Animals

Mice used in the study were housed in the Arkansas Children’s Research Institute Animal Facility. All animal studies were approved by and performed in accordance with the guidelines of the Institutional Animal Care and Use Committee of the University of Arkansas for Medical Sciences/Arkansas Children’s Research Institute. For surgeries, mice were anesthetized by inhalation of 1–5% isoflurane. Euthanasia was performed with CO₂ inhalation followed by cervical dislocation. The mouse strain CD4CreER^TM^ was purchased from Jackson Laboratories (#022356). Animals are housed at 72°F, with monthly sentinel testing conducted in each room to monitor for potential pathogens.

### T cell isolation

Single-cell suspensions enriched for naïve CD4^+^ or CD8^+^ T cells from mouse spleens were isolated. Briefly, spleens were minced and passed through a 70-μm-nylon cell strainer into serum containing PBS. After washing in PBS and lysing red blood cells, T cells were isolated from mouse splenocytes using EasySep™ Naïve CD4^+^ T or CD8^+^ T cell isolation kits (StemCell: Catalog #19765 and 19858, respectively) according to the manufacturer’s protocol. Human PBMCs were treated similarly, but with EasySep™ Human Naïve Pan T Cell Isolation Kit (StemCell: Catalog # 17961).

### Flow cytometry

Isolated CD4^+^ and CD8^+^ T cells were stained after isolated and checked for isolation specific purity prior to assay. Cells were stained for cell surface markers and resuspended in eBioscience flow cytometry staining buffer (Invitrogen, cat #00-4222-26). Viability was assessed by staining with 7-AAD or propidium iodide Viability Staining Solution. For proliferation analysis, isolated T cells were stained with cell trace violet (Invitrogen catalog#: C34557). Cell Trace violet fluorescent intensity was measured at day 0 and day 5. For apoptosis analysis, cells were stained with Annexin V and propidium iodine according to manufacturer protocol. Double positive cells were quantified for comparison.

### Metabolic analysis

Metabolic analysis was completed as described previously, specifically extracellular acidification rate (ECAR) and oxygen consumption rate (OCR) was measured using the Seahorse XFe bioanalyzer. 2 × 10^5^ CD4^+^ or CD8^+^ T cells per well (≥ 8 wells per sample) were spun onto Cell-Tak (Corning)–coated seahorse 96-well plates and preincubated at 37°C for approximately 20 min in the absence of CO2. ECAR was measured under basal conditions and in response to activation (PMA/Ionomycin), 2 μmol L−1 oligomycin and 10 mmol L−1 2-Deoxy-D-glucose (2-DG).

### Mixed lymphocyte reaction

Donor splenocytes were collected from 8 week-old BalbC mice (Jackson laboratories). Spleens were meshed through 70 µm filter, red blood cells were lysed and whole splenocytes were resuspended in supplemented RPMI. Donor splenocytes were treated with Mitomycin C (25 µg/mL) for 30 minutes prior to coculture with CD4^+^ or CD8+ T cells. Donor splenocytes were plated at a 4:1 ratio to responder splenocytes and incubated for 48 hours. Following incubation, cells were spun down and supernatant was collected for ELISA analysis of IL2 (Biolegend) and IFNγ (R&D systems) according to manufacturer protocol.

### Tumor Study

Tumor growth studies were conducted in equal numbers of male and female C57BL/6 Cre⁺-PKcs^wt/wt^, Cre^+^CD4-PKcs^fl/fl^, or Cre^+^CD8-PKcs^fl/fl^ mice. Mice were treated with tamoxifen for 7 days to induce recombination. MC-38 or B16-F10 cells were harvested during the logarithmic growth phase, washed in PBS, and implanted subcutaneously into the flank of each mouse using a 27-gauge needle (1×10^6^ cells in 100 µl) 2 days post tamoxifen treatment. Mice were monitored daily for signs of distress and tumor growth. Tumor growth was measured on alternate days using digital calipers. Tumor endpoint was established at a diameter of 20mm or earlier if mice exhibited signs of discomfort. Tumor volume was calculated using the formula: Tumor Volume=(Length×Width^2^)/2.

### Skin Transplants

Donor Balb/C tail skin was transplanted onto the backs of Balb/C mice or tamoxifen-treated male or female C57BL/6 Cre⁺-PKcs^wt/wt^, Cre^+^CD4-PKcs^fl/fl^, or Cre^+^CD8-PKcs^fl/fl^ mice, and graft rejection monitored over time. Skin graft surgeries were completed as reported previously. Prior to surgery, mice were treated with tamoxifen for 7 days, rested for one day, then underwent skin transplant. Briefly, skin grafts were harvested from the tails of 8-week-old BalbC mice from Jackson laboratories. Tails were removed, longitudinally cut and peeled from underlying tissue. Tail skin was placed in PBS on ice and cut to size, ∼200mm^3^. Recipient C57/Bl6 mice were anesthetized to prepare graft bed on the dorsal surface between front paw shoulder blades about 1 cm caudal to neck fold. Graft bed was shaved and cleaned with ethanol and iodine. Graft bed was marked with surgical pen and incised with scissors. Full thickness native skin was removed down to panniculus carnosus, a thin layer of striated muscle deep to the dermis. Grafts were then placed on graft bed and attached with 6-0 prolene monofilament non-absorbable suture (Ethicon, cat: 8706). Following attachment of graft, mice were given meloxicam for pain and removed from anesthesia. Mice were woken up on heating pad and then placed in single housed cages. Animals were treated with NU7441 (10 mg/kg in 40% PEG400/saline) or vehicle (40% PEG400/saline/DMSO) daily via intraperitoneal injection. Graft necrosis was tracked daily for 30 days after surgery. Animals were euthanized upon graft death (>80% necrosis).

### Statistical analysis

The data in this manuscript were analyzed using GraphPad Prism 7. A significance threshold of α = 0.05 was applied, with p ≤ 0.05 considered statistically significant. Details on the number of experimental replicates, animals per experiment, and statistical tests used are provided in the respective figure legends.

