## Supplementary figures and images for "DNA-PKcs governs LAT-dependent signaling in CD4^+^ and CD8^+^ T cells"

### Figure S1 (1).tiff

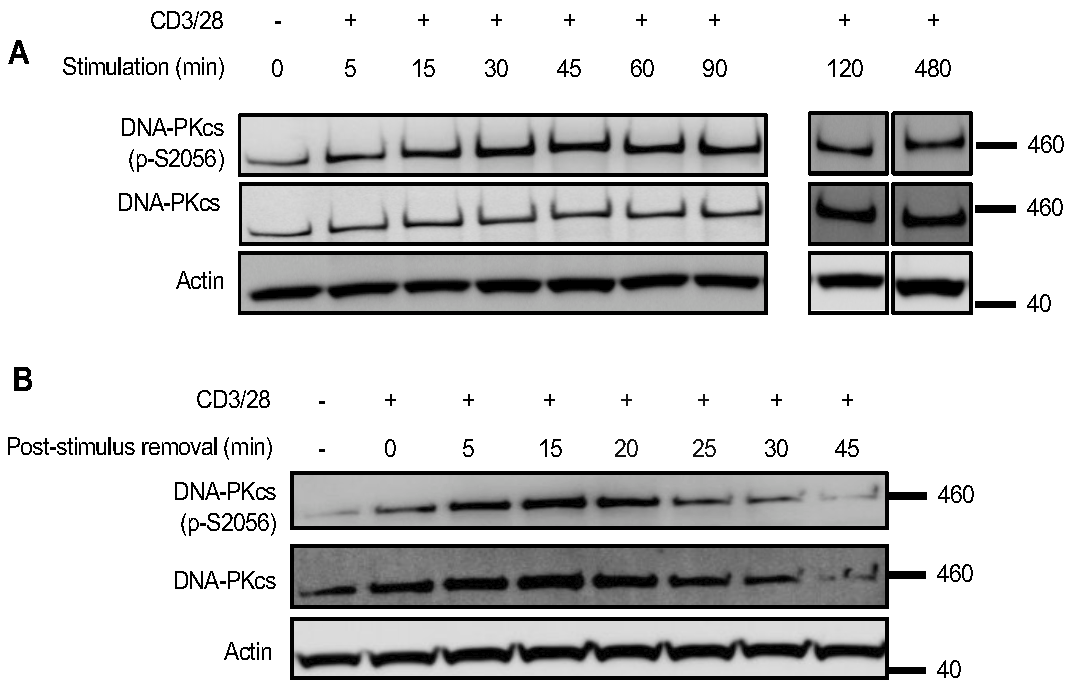

### Figure S2.tiff

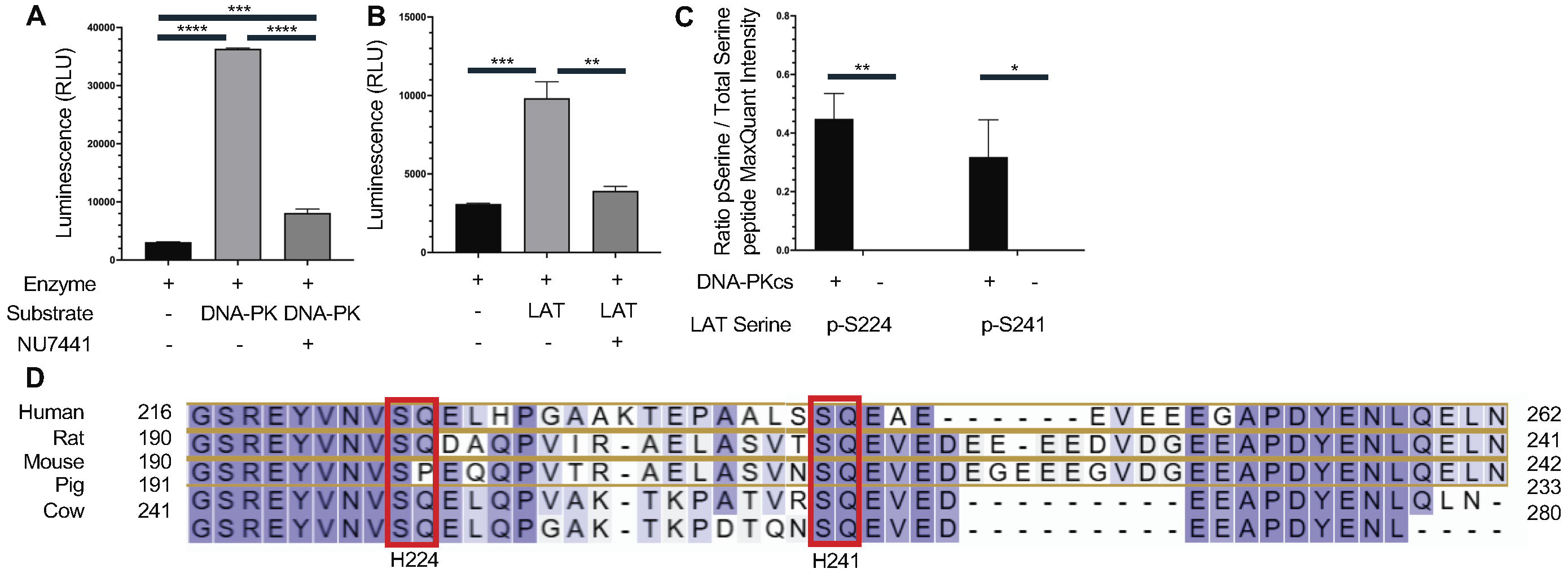

### Figure S3.tiff

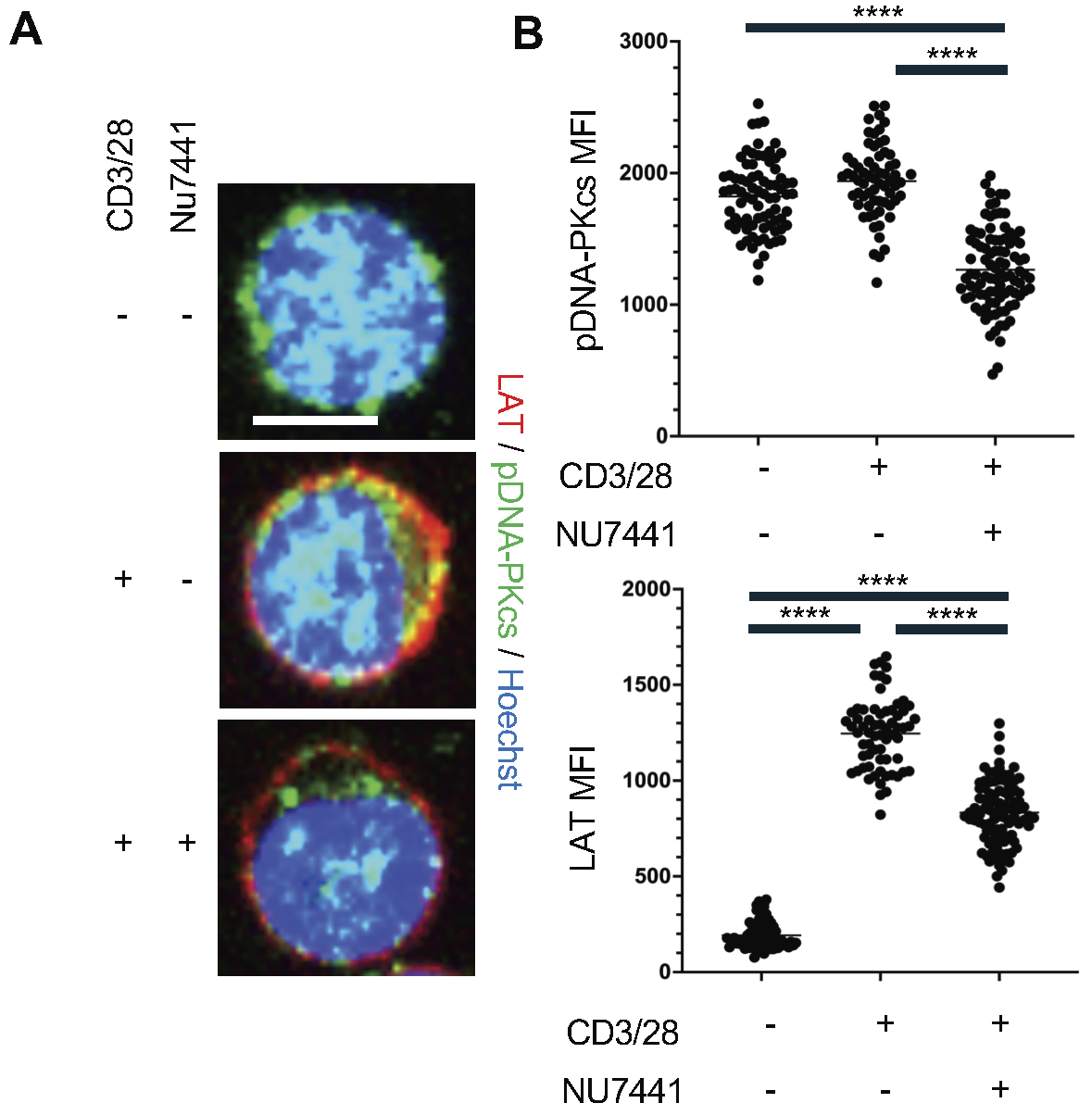

### Figure S4.tiff

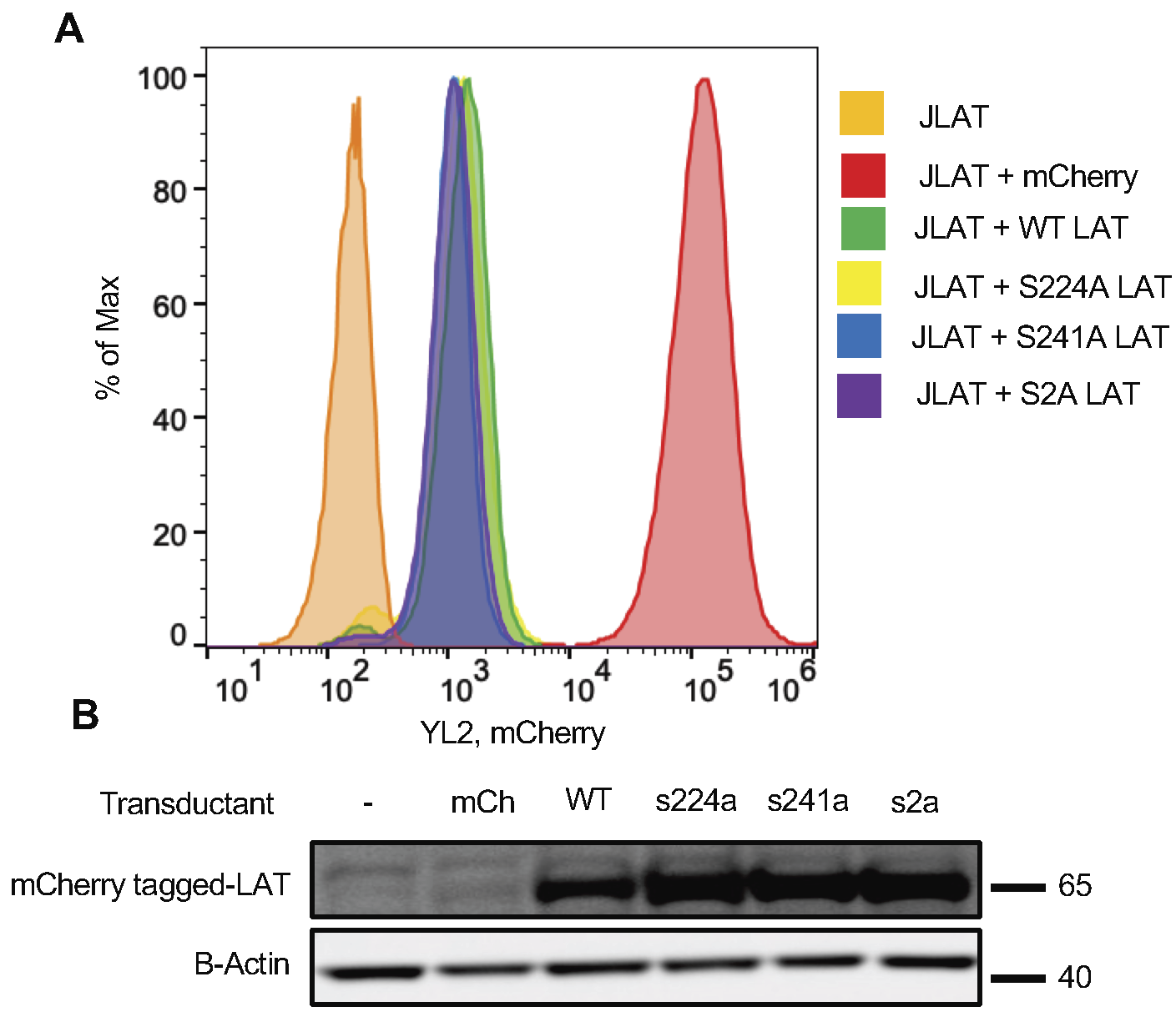

### Figure S5.tiff

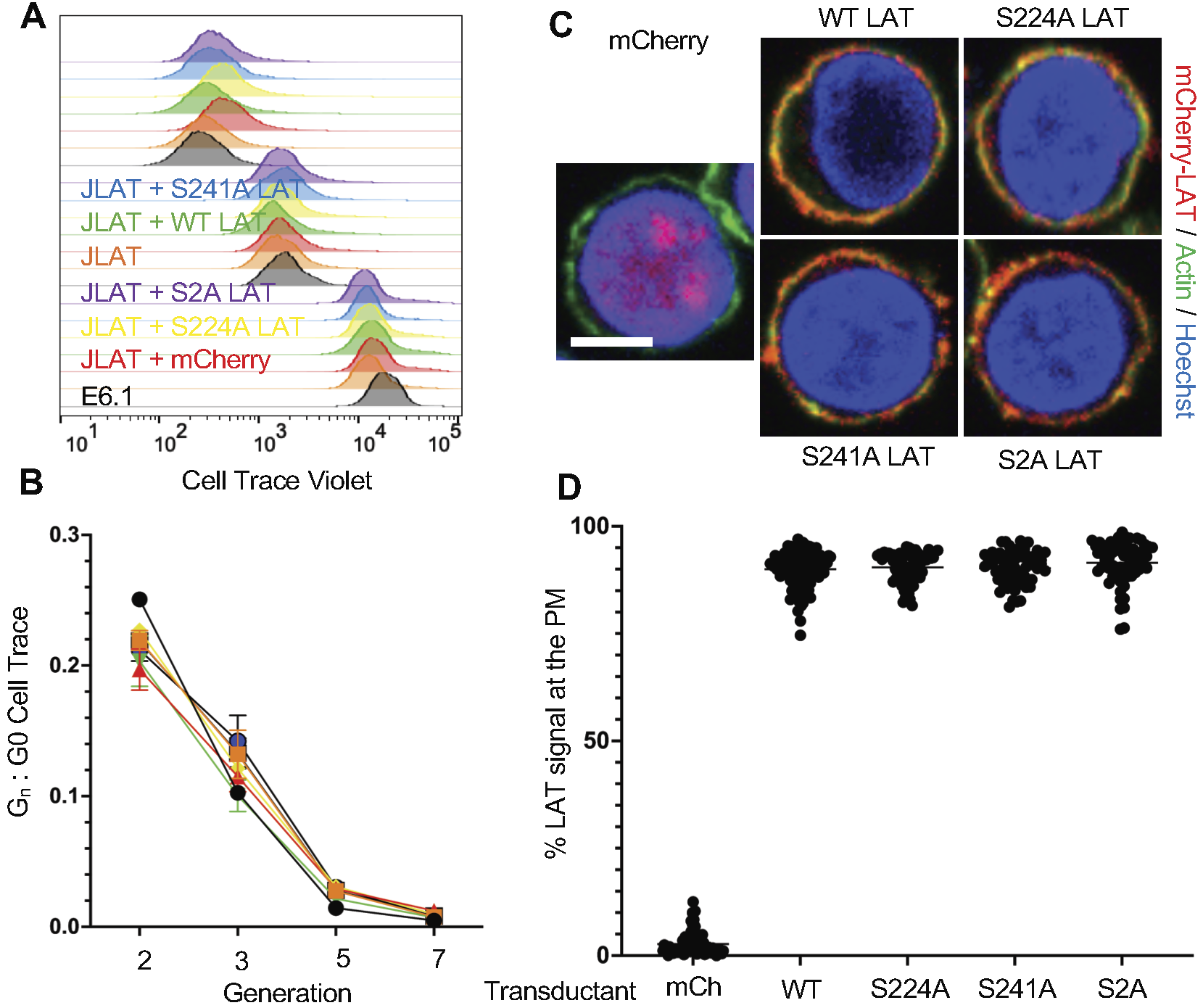

### Figure S6.tiff

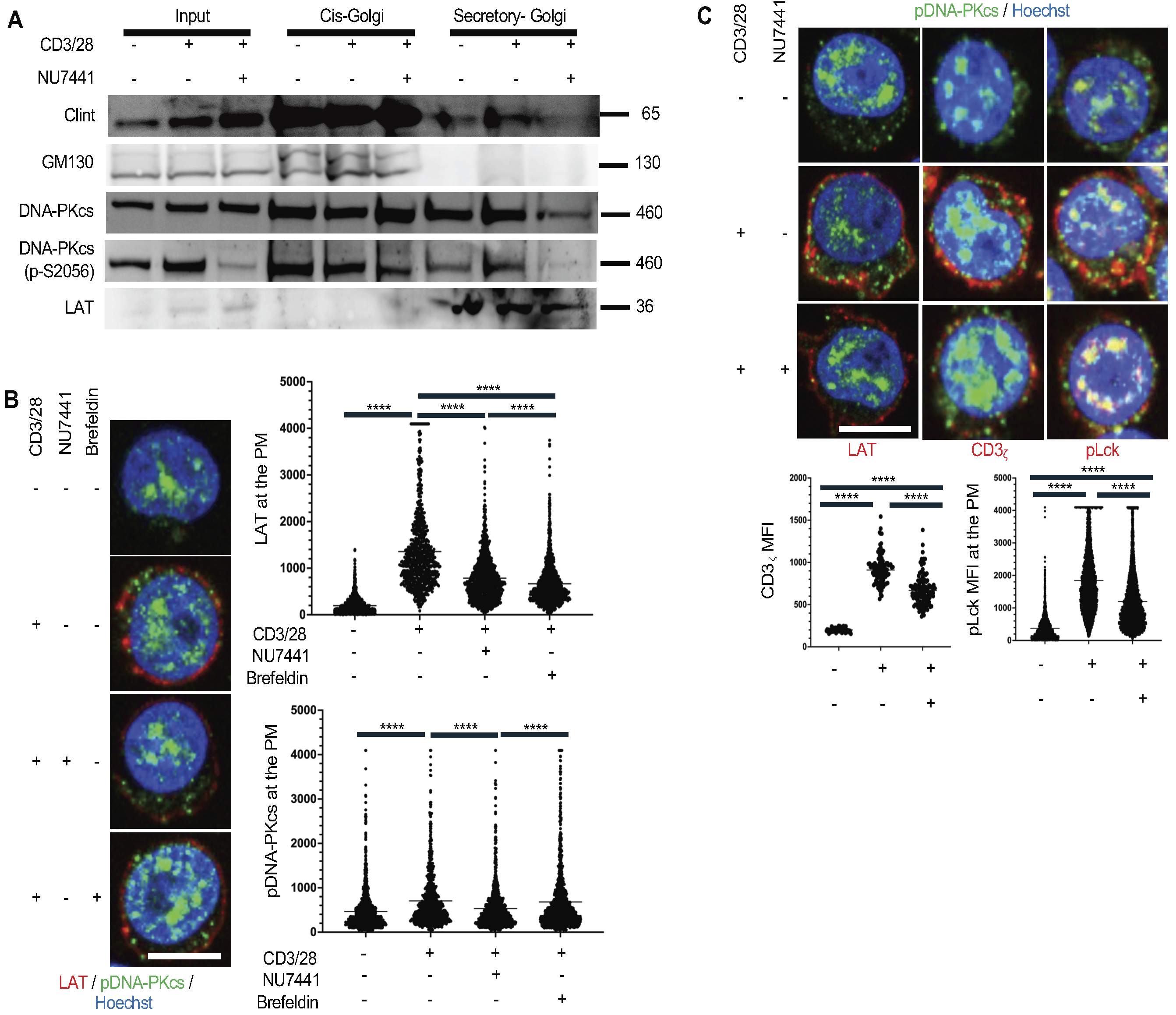

### Figure S8.tiff

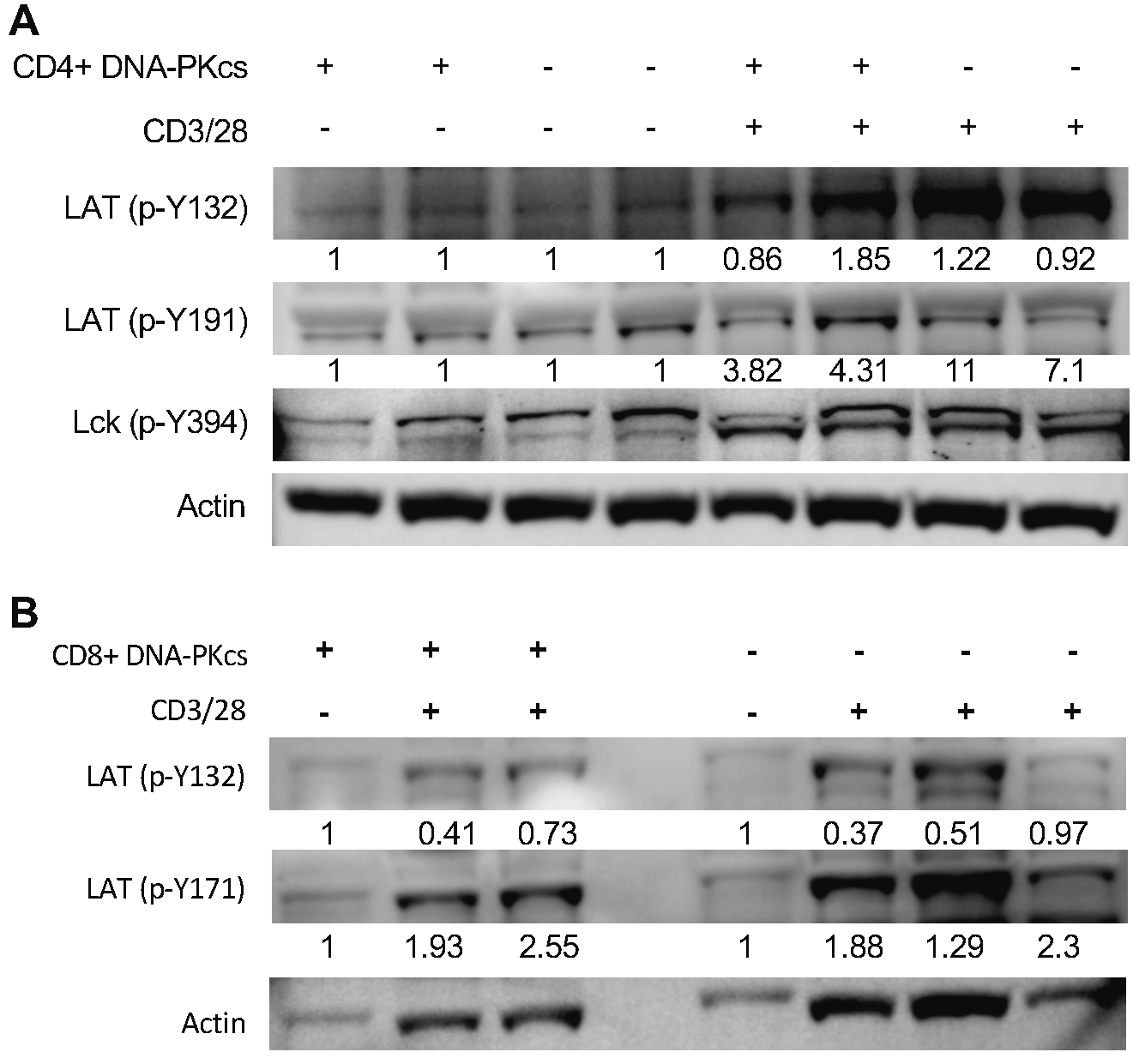
